# Macroscale Connectivity in the Octopus brain

**DOI:** 10.1101/2025.05.28.656524

**Authors:** Federica Pizzulli, Marisa Barjami, David B. Edelman, Graziano Fiorito, Giovanna Ponte

**Author notes:** These Authors mutually contributed (Federica Pizzulli, Marisa Barjami). **Corresponding Authors**: Federica **Pizzulli** Department of Biology and Evolution of Marine Organisms Stazione Zoologica Anton Dohrn Napoli, Italy; Giovanna **Ponte** Department of Biology and Evolution of Marine Organisms Stazione Zoologica Anton Dohrn Napoli, Italy.

## Abstract

Comparative studies support the existence of functional analogies between given areas in the octopus and mammalian brains. Despite marked phylogenetic distance, the central nervous system of cephalopod mollusks is characterized by a complex network of anatomically interconnected and interacting neuronal populations. Previous studies provide a thorough description of the octopus neural network through neuronal tracing of axonal projections, offering a unique opportunity to systematically examine the brain’s network architecture. Here, we chart the topological organization of *Octopus vulgaris* neural network including information on the macroscale interregional pathways between 32 cortical and subcortical regions as provided by a high-quality historical dataset. We found more than 350 main neural connections (afferent, efferent) encompassing the three masses of octopus brain (supra-, sub-oesophageal masses, optic lobes). The octopus brain network possesses multiple nonrandom features promoting segregation and integration, including near-minimal path length, multiscale community structure, hubs, small-world features, and motif composition. Taken together, these attributes support an association between network architecture and function and are consistent with studies in a range of other species, suggesting the existence of a set of universal organizational principles across phylogeny. These findings expand our understanding of neuronal structures by highlighting brain regions that had previously received less attention.

## Introduction

Studies on the structure and architecture of nervous systems of various species reveal the existence of common organizational principles of networks supporting sophisticated neural and behavioral repertoires. A comparative outlook at brain network properties shows shared neural phenotypes, cross-species analogies (and/or homologies), and the emergence of complex brain structures and functions. Despite the differences in scale and complexity, some fundamental principles of neural organization appear to be conserved across vertebrates and invertebrates (e.g., Ardesch et al., 2019; Kaiser & Varier, 2011; Lin et al., 2024; Martinez & Sprecher, 2020; Nieder, 2023; Towlson et al., 2013).

The topological structure, architecture, and computational ability of a brain is intimately linked to function via the overall and regional connectivity of its network and the various potential pathways for signal integration (e.g., Avena-Koenigsberger et al., 2018; Hagmann, 2005; Leergaard et al., 2012; Seguin et al., 2023). The analysis of neural networks (connectomes) shows abilities to: **a.** process specialized information, and **b.** integrate neural information across different domains. Among various features, connectomes are recognized for local densely clustered communities, ensuring segregation of information and local specialization, global shortcuts providing an infrastructure for communication between remote regions. Through network analysis, the wiring of neural networks revealed that brain networks in different species display cost-effective wiring and a small-world modular organization with high levels of local clustering and modular structure, combined with short communication pathways ensuring global relay, and functionally specialized modules intertwined by highly connected central hubs (e.g., Bullmore & Sporns, 2012; Seguin et al., 2023).

Cephalopod brains are recognized as complex neural structures that largely diverged from the ganglionic arrangement of their molluscan origin (2015; Shigeno et al., 2010). These ‘brains’ allow sophisticated behavioral repertoires and support a wide range of learning and memory processes, cognition, and possibly consciousness (Marini et al., 2017; Nixon & Young, 2003; 2022; Ponte et al., 2021; Wang & Ragsdale, 2019). Despite decades of research highlighting their neural complexity, sophisticated behavioral repertoire, and cognitive abilities, the structural features of cephalopod brains—such as inter-areal circuitry and large-scale functional connectivity—have been explored in only a limited number of studies.

Among cephalopods, the brain of *Octopus vulgaris* is known for its relative size (i.e. brain-to-body mass ratio), the large number of nerve cells (about 200 million), the possession of the most centralized (or ‘cephalized’) nervous system among all extant cephalopods, thus rivalling higher vertebrates (Kandel, 1979; Packard, 1972; Young, 1963). An alternative among invertebrates is the ’mini-brain’ of the honeybee, which presents an intriguing comparative case. It contains approximately 960,000 neurons densely packed into a volume of one cubic millimeter, accounting for a rich repertoire of complex behaviors (Menzel & Giurfa, 2001; Menzel et al., 2006).

In addition to the high degree of centralization, cephalopod brains are characterized by ***i.*** large populations of very small neurons, estimated more than 50% of the overall neural cell population, with nuclear sizes ranging from 3-5µm, in given regions, ***ii.*** functional analogies with mammalian brain areas of numerous octopus’ brain structures, ***iii.*** compound field potentials like those identified in vertebrate brains, ***iv.*** a rich diversity of neuro-transmitters/neuromodulators and a characteristic molecular fingerprint including the existence of brain-specific orphan genes and retrotransposons, to mention some (e.g., Fiorito et al., 2020; Petrosino et al., 2022; Ponte & Fiorito, 2015; Ponte et al., 2021; Styfhals et al., 2019; 2022).

*O. vulgaris* is perhaps the most famous and best studied of all octopus species (review in e.g., Marini et al., 2017; Sanders, 1975; Wells, 1978; Zarrella et al., 2019). Since the late 1940s, J.Z. Young and a plethora of collaborators working at the Stazione Zoologica of Napoli (Italy) carried out a systematic analysis of the neural structures underlying behavioral plasticity in this animal (e.g., Boycott, 1954; Boycott & Young, 1955; Maldonado, 1963b; Packard, 1963; Young, 1960). Based on these studies John Zachary Young (JZ) provided an accurate and detailed description of the gross morphological features, organization, architecture (e.g., inter-areal connectivity), and neural arrangement of the brain of *O. vulgaris*, featuring a *model of the brain* (Young, 1964; 1971).

The *corpus* of the monumental work of JZ has been based on both a vast collection of histological sections from octopus brains and other nervous structures, and a series of experiments carried out to explore the functional division of labor among, or co-participation of, the different areas of the central brain and other regions of the octopus’ nervous system in modulating *O. vulgaris* behavioral responses and learning abilities (e.g., Maldonado, 1963b, 1965; Marini et al., 2017; Sanders, 1975; Young, 1991, 1995). In the original work cellular composition of, and relationships between, brain regions (i.e. lobes and parts) have largely been based on Golgi and reduced silver techniques. In addition, JZ provided a detailed account of tracts, connectives, commissures, nerves, and the network of fibers (Young, 1932; 1963, 1971).

Here we analyze the macroscale connectivity of *O. vulgaris* brain (JZ data: Young, 1971). We then investigated a range of topological attributes and brain network features, including highly connected hubs, community structure, motif composition, rich club organization, and functionally specialized modules.

## Results

Based on the identification of neural pathways (afferent and efferent, n= 439) linking different areas of *O. vulgaris* brain the connectivity matrix (Figure 1; see also Supplementary Info and Supplementary Figure 2) includes 249 directed edges between the 32 cortical and subcortical regions (nodes). The number of directed links here accounted is underestimated: we did not include in this analysis paths or connections reaching the peripheral nervous system or targets external to the brain (e.g., ophthalmic and oculomotor nerves, mandibular- radular and salivary nerves, visceral and cardiac nerves; see also Supplementary Table 1). For further details about network reconstruction and identification of areas in the octopus’ brain see Supplementary Info. The number of inward links (249) counts for slightly different values when outward edges (245) are considered; however, inward and outward edges achieve comparable values when considering the entire set. The number of edges reflect the existence of an anatomical projection between brain regions; therefore, we assume a slightly biased connectivity (see below). It is noteworthy to mention that we were unable to provide any information on the level of connectivity strength of edges. Our dataset did include, only in a few instances, information on the number of fibers for a given tract or neural pathway in the neuropile (see Supplementary Table 3); therefore, we preferred to avoid any estimation of edge strength to avoid potential bias.

**Figure 1.**
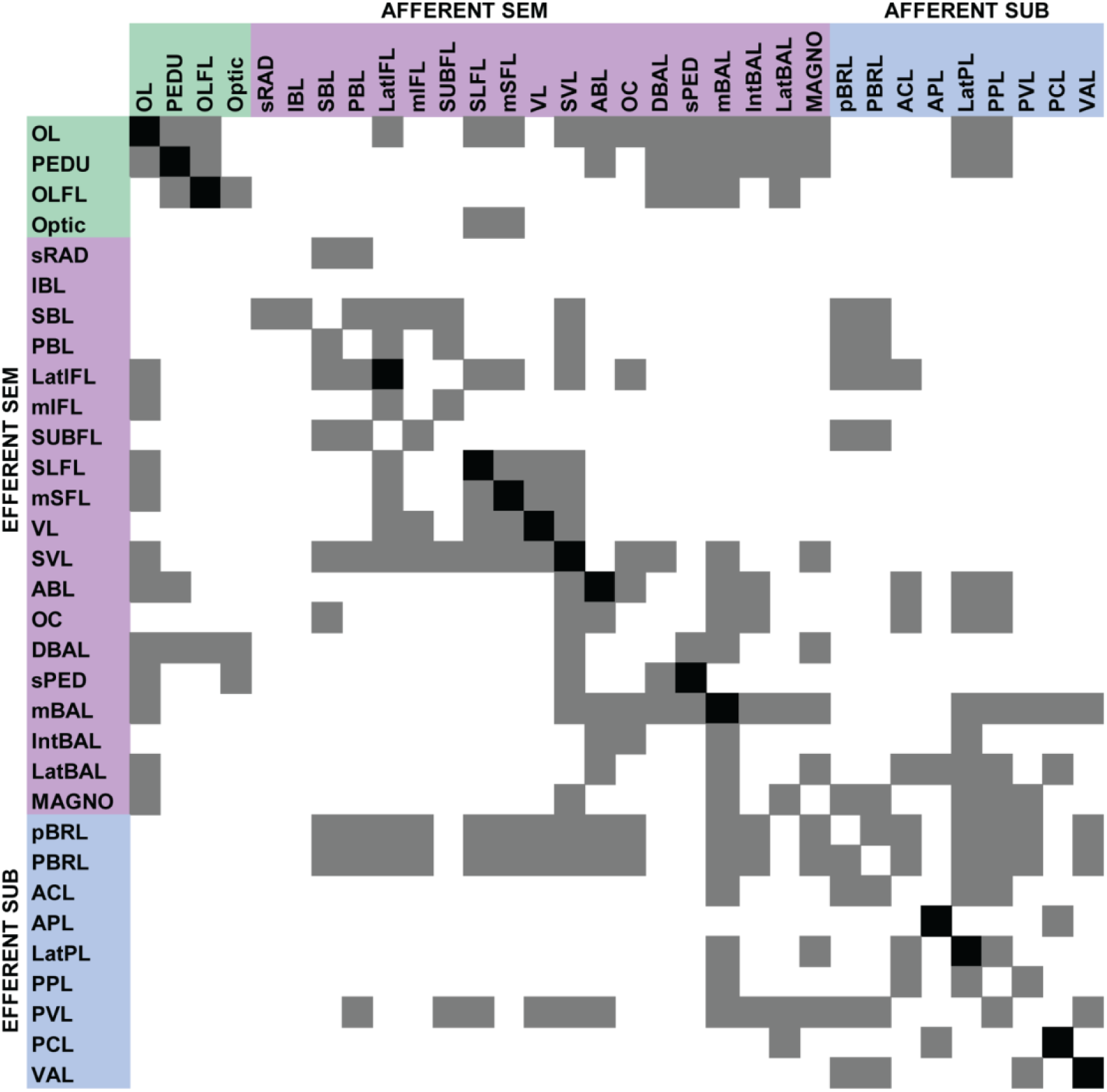
Connectivity matrix of the *O. vulgaris* neural network based on J.Z. Young dataset (1971< see Supplementary Info). The binary matrix accounts for afferent and efferent connections noted between the brain lobes identified within the supraoesophageal (SEM), suboesophageal (SUB) masses and optic lobes (OL). A directed binary adjacent matrix of size N × N (N = 32) was constructed (Supplementary Figure 2) expressing the presence of an anatomical projection between two regions (coded as 1, here in black) and absent and/or non-reported connections coded as a 0 (here in white).

Table 1 summarizes the in-degree (id), out-degree (od) and nodal degree (nod) representing the total number of links connected to a given node. In *O. vulgaris* brain network, the most connected regions resulted to be the medial basal lobe (mBAL: id = 17, od = 15, nod = 32), the optic lobes (OL: id = 13, od = 17, nod = 30), the subvertical lobe (SVL: id = 17, od = 14, nod = 31) and brachial lobes (id = 9, od = 19, nod = 28; same degree values for: prebrachial, pBRL; postbrachial, PBRL). The three regions (medial basal, subvertical and optic lobes) also have the greatest indegree (17, 17 and 13 respectively) together with the lateral pedal and post-pedal lobes (id = 13 and 12). As mentioned, the prebrachial (pBRL) and the postbrachial (PBRL) lobes count for the higher number of outgoing connections (od = 19), followed by the optic lobes, medial basal, subvertical and the palliovisceral lobe (od = 14; see Table 1). Overall, eighteen octopus brain regions have more afferences than efferences (belonging to SEM: 10 nodes, SUB: 7 nodes; see also Supplementary Info), ten (10) nodes show the opposite trend (outdegree > indegree; belonging to SEM: 6, SUB: 3), while in four cases *O. vulgaris* brain areas have the same number of incoming and outgoing connections (all belonging to the supraoesophageal mass: Lateral inferior frontal, subfrontal, anterior basal, precommissural lobes).

**Table 1.**
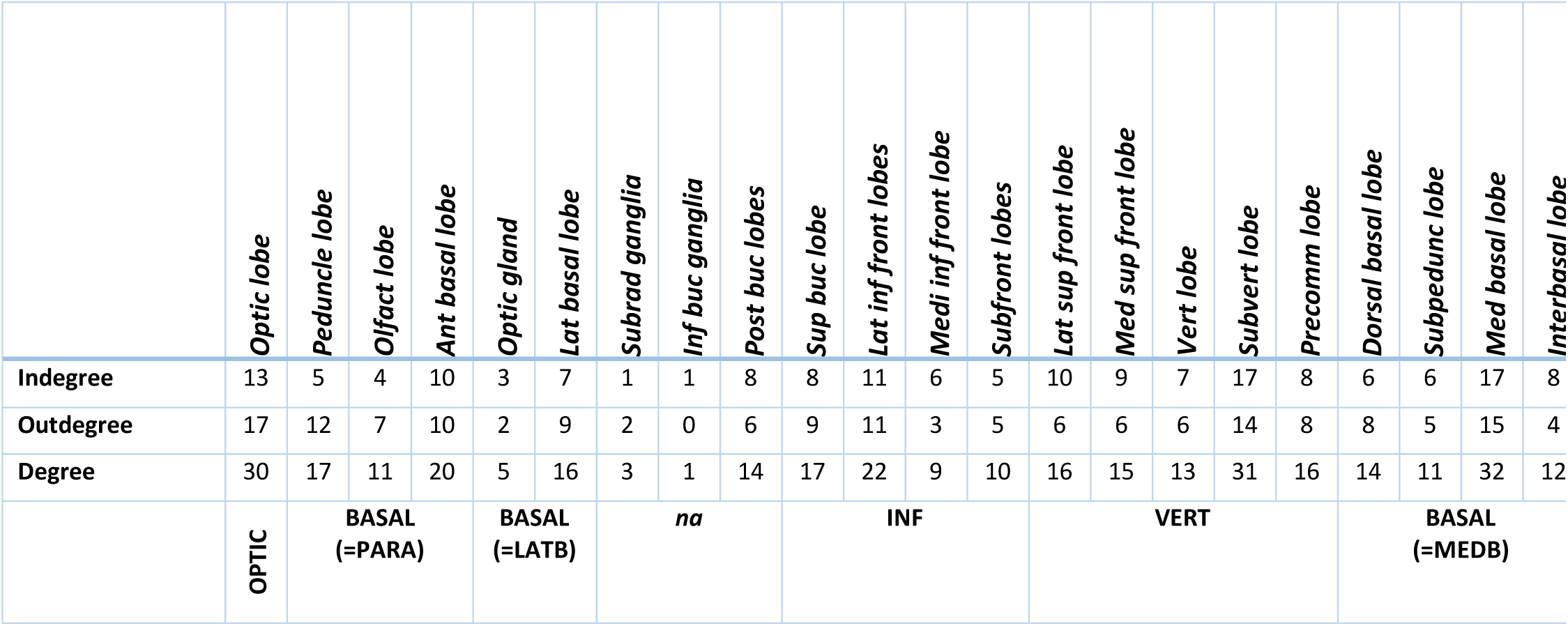
Indegree (total number of afferent connections), outdegree (total number of efferent connections), and degree (total number of afferent and efferent connections) for lobes pertaining to optic lobe and supra- and suboesophageal mass of *O. vulgaris* brain. The last row groups lobes into functional sets as identified by Maddock and Young (1987); see text for details.

|  | *Optic lobe* | *Peduncle lobe* | *Olfact lobe* | *Ant basal lobe* | *Optic gland* | *Lat basal lobe* | *Subrad ganglia* | *Inf buc ganglia* | *Post buc lobes* | *Sup buc lobe* | *Lat inf front lobes* | *Medi inf front lobe* | *Subfront lobes* | *Lat sup front lobe* | *Med sup front lobe* | *Vert lobe* | *Subvert lobe* | *Precomm lobe* | *Dorsal basal lobe* | *Subpedunc lobe* | *Med basal lobe* | *Interbasal lobe* |
| --- | --- | --- | --- | --- | --- | --- | --- | --- | --- | --- | --- | --- | --- | --- | --- | --- | --- | --- | --- | --- | --- | --- |
| Indegree | 13 | 5 | 4 | 10 | 3 | 7 | 1 | 1 | 8 | 8 | 11 | 6 | 5 | 10 | 9 | 7 | 17 | 8 | 6 | 6 | 17 | 8 |
| Outdegree | 17 | 12 | 7 | 10 | 2 | 9 | 2 | 0 | 6 | 9 | 11 | 3 | 5 | 6 | 6 | 6 | 14 | 8 | 8 | 5 | 15 | 4 |
| Degree | 30 | 17 | 11 | 20 | 5 | 16 | 3 | 1 | 14 | 17 | 22 | 9 | 10 | 16 | 15 | 13 | 31 | 16 | 14 | 11 | 32 | 12 |
|  | OPTIC | BASAL<br>(=PARA) |  |  | BASAL<br>(=LATB) |  | na |  |  | INF |  |  |  | VERT |  |  |  | BASAL<br>(=MEDB) |  |  |  |  |

As mentioned above, the J.Z. dataset did not systematically report the number of fibers for afferent and/or efferent tracts/connections (see also Supplementary Info). Consequently, we were unable to estimate the weight of each degree. However, a given degree may represent thousands of fibers (see Discussion for a rough estimation as an example).

The reconstructed *O. vulgaris* connectome revealed a density of 25.1%.

In random networks all connections are equally probable resulting in a symmetrically centered degree distribution (Gaussian). In contrast, complex networks show non-Gaussian degree distributions, often with a long tail towards high degrees. In the brain network of *O. vulgaris* in-degree and out-degree distributions are right-skewed (Supplementary Figure 3), confirming that the probability of a high degree node is greater than would be expected in an equivalent random network, but less than that in a scale-free network.

Based on our data, 64.3% of all connections were found to be bidirectional, and 35.7% consisted of unidirectional pathways. Considering the functional sets identified by Young and coworkers (see also Fig. 1) as an indicator of brain modularity, bidirectionality was recognized in 56 % of all intermodular connections (44 % thus being unidirectional). In contrast, the majority of intramodular connections (75 %) were found to be bidirectional (25 % being unidirectional).

### Centrality and ‘Small World’ Architecture

The assortativity of *O. vulgaris* network is positive (0.036; see Figure 2) and indistinguishable from the null distribution, suggesting no evidence of homophilic attachment among nodes with similar strengths. Octopus brain network results to be about 1.7 times above random levels of clustering (Figure 2; CC = 0.64; normalized clustering coefficient γ = 1.7) with short characteristic path length (characteristic path length = 2.12; normalized characteristic path length λ = 1.12; Fig. 2). The above characteristics support the existence of a small world property (small-world index σ = 1.52).

**Figure 2.**
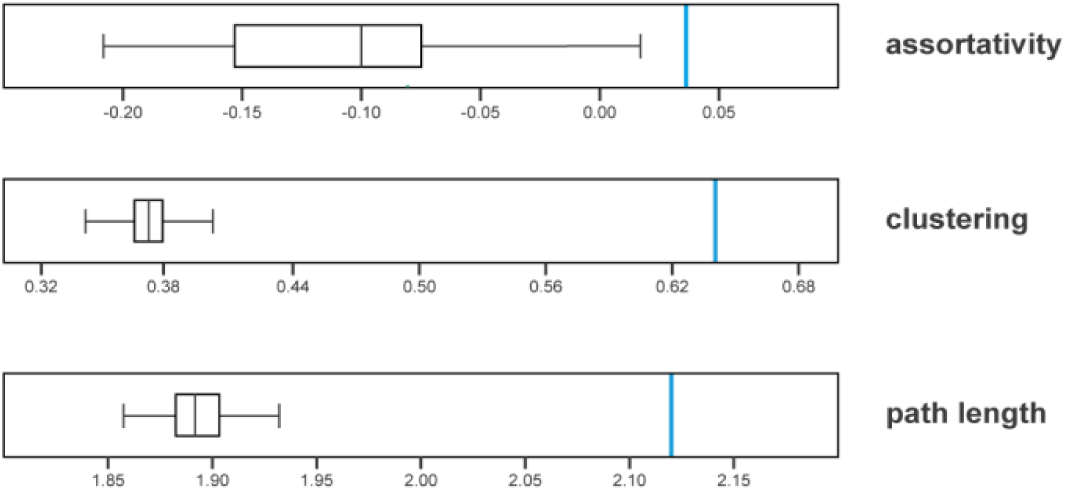
Global topological attributes of the octopus brain network. Values for Assortativity – i.e. tendency for nodes to link with similar nodes; clustering (proportion of completed triangles around a node), and path length, as mean shortest set of edges connecting all pairs of nodes in the network are). *O. vulgaris* network measures are shown in light blue; distributions from 1,000 in-degree and out-degree preserving rewired null networks are shown in gray. See text for details.

Further examination of global organization shows that the network possesses a global efficiency (E*_glob_* = 0.577, normalized E*_glob_*= 0.969) corresponding to about 0.97 times that of randomized networks of equal density and equal degree sequence, reflecting a relatively globally oriented organization. In addition, local efficiency is 1.15 times higher than the randomized network (E*_loc_* = 0.71; normalized E*_loc_* = 1.14). High efficiency of parallel information transmission is achieved in only about 25% of total possible connections (connection density = 0.251), reflecting an optimization of wiring cost in the brain, i.e., a high efficiency achieved with a reduced number of anatomical connections.

### Community structure and modularity

We next attempted to characterize the macroscale structure of the *O. vulgaris* brain network by identifying architectural properties dictating the organization of neural elements into circuits and clusters. Following Rubinov and Sporns (2010), we applied a Louvain-like greedy modularity maximization approach in order to identify multiscale communities, and performed a parameter search across a range of resolutions to explore the possible existence and composition of communities from low to high resolution (i.e. small set of large communities, vs a larger set of small communities; Figure 3). The community structure of the network (i.e. its modularity) represents subdivisions into non-overlapping groups of nodes, maximizing the number of within-group links, while minimizing the number of between-group links (Girvan & Newman, 2002).

**Figure 3.**
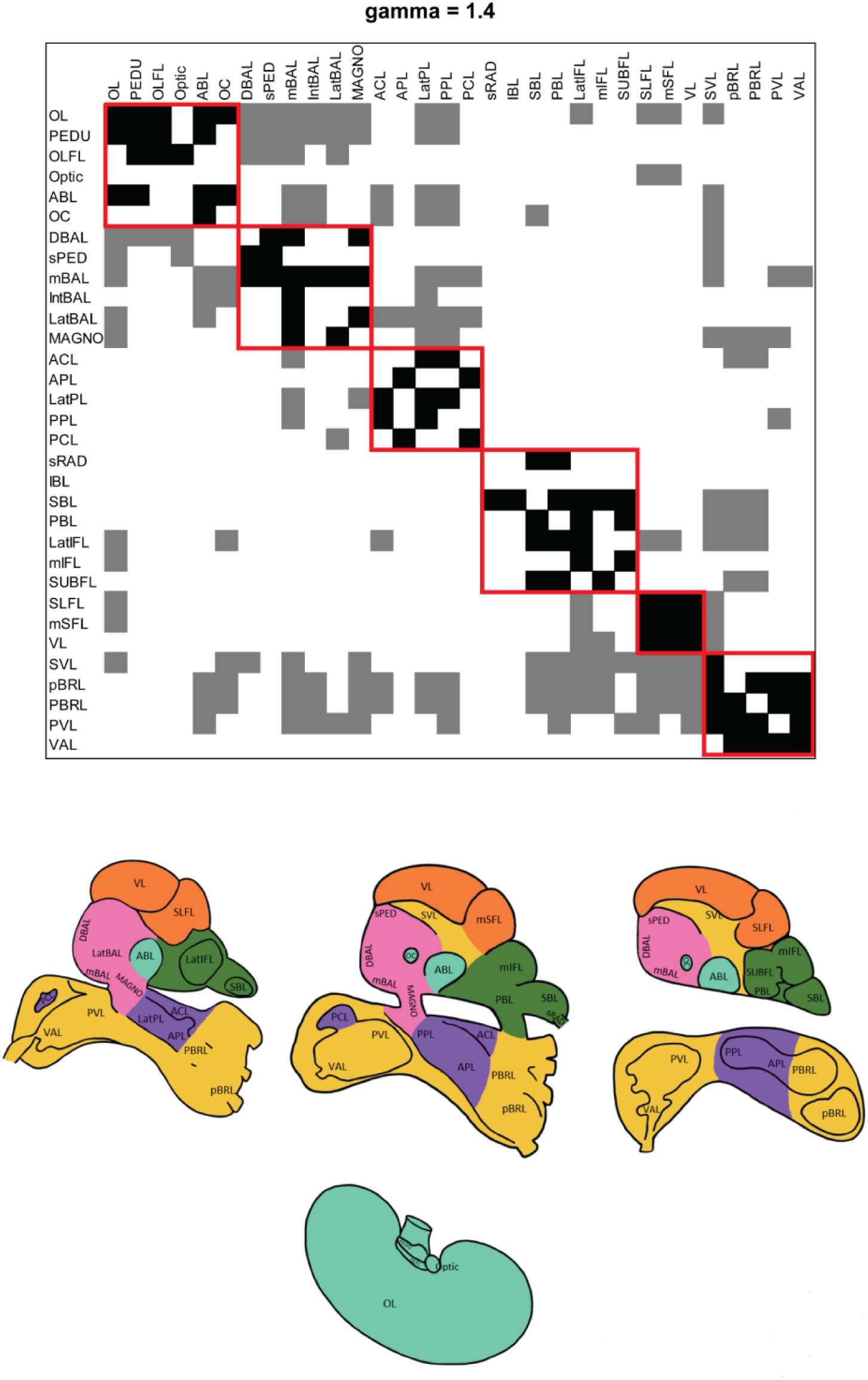
Community structure detected in the brain network of *O. vulgaris*. The adjacency matrix is reordered by community assignment (top); the flatmap projections on sagittal and parasagittal sections of octopus brain and its parts depicting the community assignments. With ***γ=1.4*** six communities were detected (red squares). The first community is represented by: optic lobe, peduncle, olfactory, anterior basal and precommissural lobes. The second by: dorsal basal, subpedunculate, median basal, interbasal, lateral basal and magnocellular lobes. The <u>third</u> including: anterior chromatophore, anterior pedal, lateral pedal, posterior pedal and posterior chromatophore lobes. The fourth community: subradular, superior buccal, posterior buccal, lateral inferior frontal, median inferior frontal and subfrontal lobes. The fifth consisting in: lateral superior frontal, median superior frontal and vertical lobes. The sixth and last one by: subvertical, pre- brachial, post-brachial, palliovisceral and vasomotor lobes.

The procedure yielded three points in parameter space with two (γ = 0.6), four (γ =1) and six (γ= 1.4) spatially contiguous communities (see red marked areas in Figure 3 and Supplementary Figure 4). At the different resolutions considered, the optic lobe and the morphologically closely related areas (peduncle-, olfactory lobes and optic gland), together with the vertical-, subvertical- and the superior frontal lobes are consistently identified as separate communities; these correspond to OPTIC and VERT functional brain-set identified by Young, respectively. At the lowest resolution, the remaining areas are distributed among the two communities: one mainly contains motor, premotor, visceral control regions and visual somatosensory cortex-like, while the other belongs to chemo- tactile and associative areas. There is no clear-cut separation of areas belonging to supra- and suboesophageal masses in the attribution to these two communities (Suppl. Figure 4). A similar pattern is found at γ =1 (Suppl. Figure 4). At the highest resolution (γ = 1.4; see Fig. 3), the six communities identified include visual sensory-motor centers, lobes belonging to the neural control of viscera and body patterning, feeding behavior, and associative centers. The analysis highlights the existence of a multiscale community structure that maps onto increasingly specialized and/or interconnected functional domains in *O. vulgaris* brain.

### Hub and Rich Club Organization

The presence of communities in the *O. vulgaris* brain network facilitates information segregation and the connectivity of specialized neural circuits; however, complementary integration remains essential. To better describe the complex balance between segregation and integration in the octopus’ network, we examined the presence of hubs and their role within the community structure. We focused on the resolution that maximizes the modularity parameter (γ = 0.6, two modules) and computed the within-module degree z-score and the participation coefficient (P; see Supplementary Table 6). Considering the indegree, five brain areas (∼15% of the total nodes) resulted with z values greater than the threshold level and can be identified as network hubs (Figure 4): the medial basal lobe (z = 2.49), the lateral pedal lobes (z = 1.77), the subvertical lobe (z = 1.41), the lateral inferior frontal (z = 1.41) and the posterior pedal lobe (z = 1.04). Among these five hubs, the lateral inferior frontal lobes (P = 0.17) and the lateral pedal lobes (P = 0.26) are detected as provincial hubs, whereas the subvertical (P = 0.48), medial basal lobe (P = 0.36) in the supraoesophageal mass, and the posterior pedal lobe in the suboesophageal mass (P = 0.38) are connector hubs. Focusing on the outgoing connections, we detected (again about 15% of the total nodes; Figure 4) four connector hubs (optic lobe: z = 1.82, P = 0.36; medial basal lobe: z = 1.56, P = 0.32; prebrachial and postbrachial lobes: z = 1.5, P = 0.49) and only one provincial hub, the peduncle lobe (z = 1.56; P = 0).

**Figure 4.**
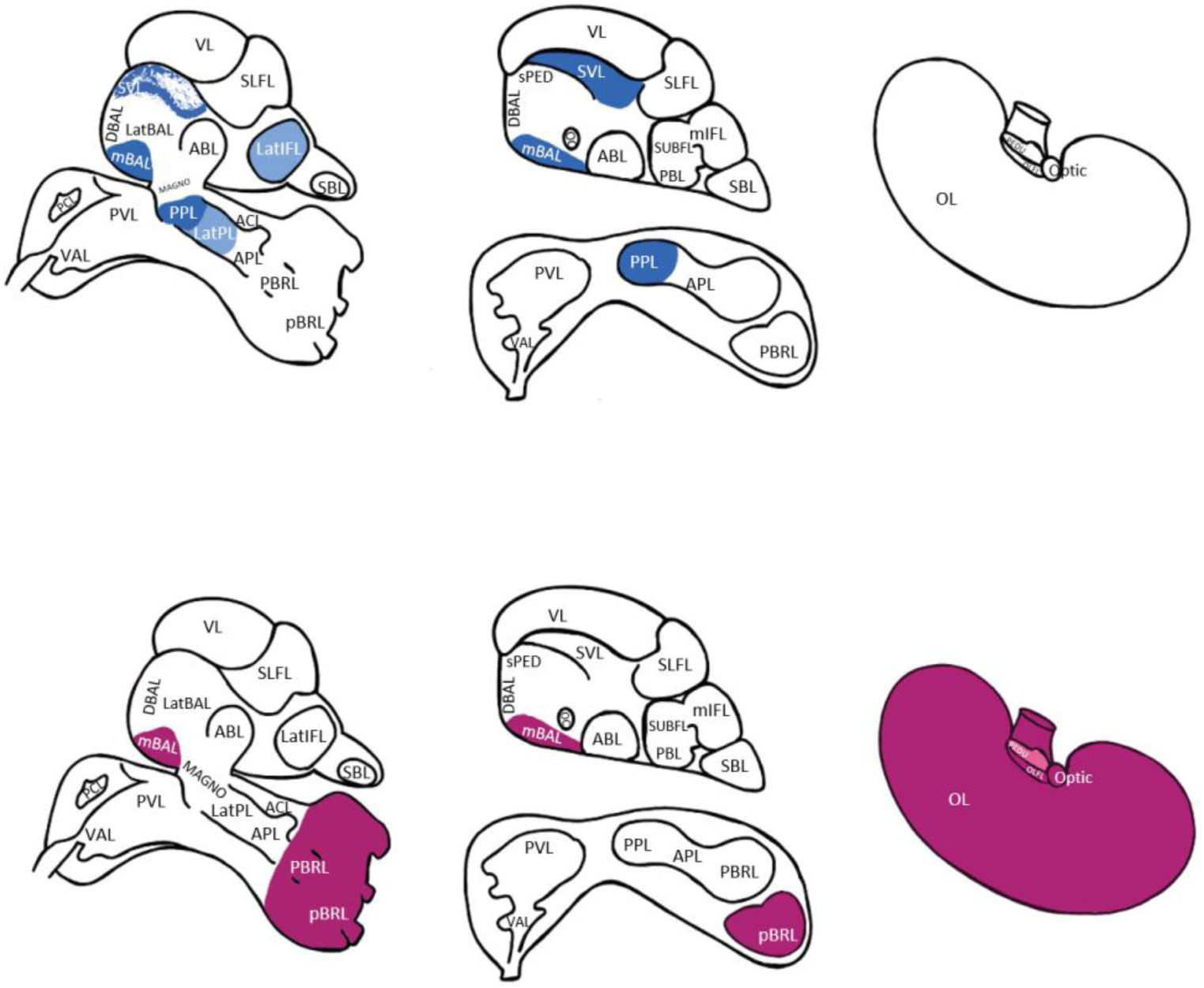
Hubs in the brain of *O. vulgaris* for indegree (blue) and outdegree (purple). The outline of the brain (supra- and suboesophageal masses) is presented at a parasagittal section (left) and sagittal (middle); optic lobes on the right. Hub nodes are marked, provincial hubs in lighter color; non-hub nodes are not colored. Note that the subvertical lobe is marked also at the ideal parasagittal section to highlight its relevance and relationship with other structures. See text for details.

The octopus neural network revealed a rich club organization. Across the selected range we identified a subset of high-degree connectivity significantly exceeding the 1,000 random networks examined (Figure 5), an observation consistent with the findings of studies on rich club organization in other species. Anatomically, a rich club organization, with a group of high-degree strongly interconnected nodes, appears evenly distributed throughout the octopus brain including the prebrachial- and postbrachial lobes (suboesophageal mass), the optic lobes, and subvertical and medial basal lobes (supraoesophageal mass). Notably, rich club nodes are evenly distributed among the segregated communities, providing an infrastructural feature to sample and integrate information from specialized domains.

**Figure 5.**
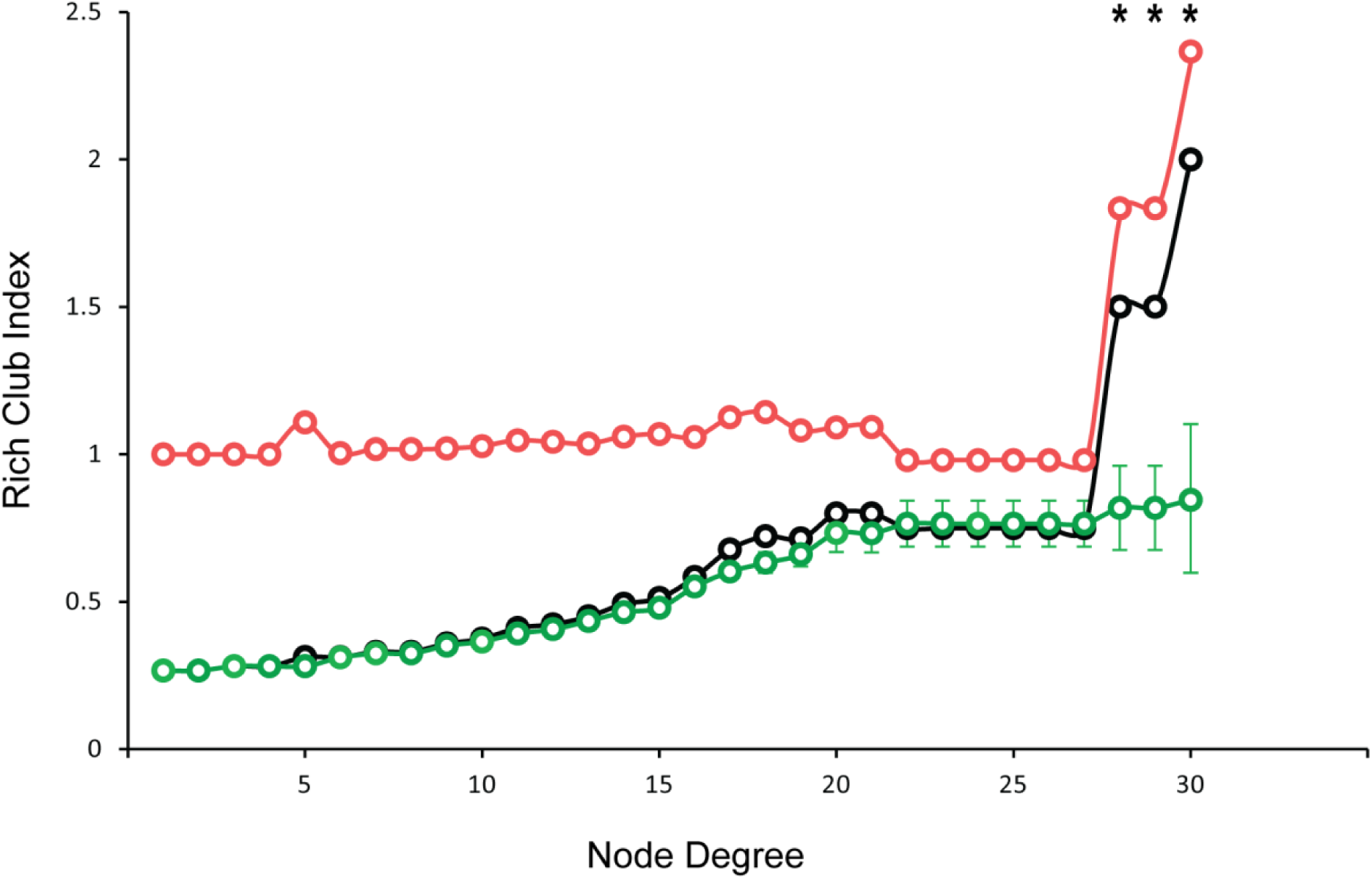
The plot shows the rich club coefficient R(k) of the brain network of *O. vulgaris* (black) for a range of k= 1 to k= 32. The mean rich club curve R random, after a set of 1000 comparable random graphs (green), is also shown and calculated to randomize the connections within the octopus brain network while preserving degree sequence. Their ratio (Normalized rich club ratio, red) is also shown. Error bars on the Rrand mark the standard deviation of of the rich club coefficient across the 1000 random networks. We identified the rich club as a subset of high-degree neurons that have a significantly greater density of connections between them than would be expected in a subset of equally high-degree nodes in a random graph.

### Motif composition of the brain network

To estimate the motif composition of the network, we focus exclusively on three-node motifs, allowing for 13 distinct configurations. Expanding to four-node motifs results in 199 possible combinations, requiring prohibitive computational analysis. We calculated network motif fingerprints - representing the frequency of motif occurrences - and compared them to an ensemble of rewired null networks (Figure 6). Comparison to the null model reveals that the *O. vulgaris* neural network is enriched for several specific motifs, including motifs 9 and 13; the octopus brain network is deficient in several motifs (1–5, 7 and 11).

**Figure 6.**
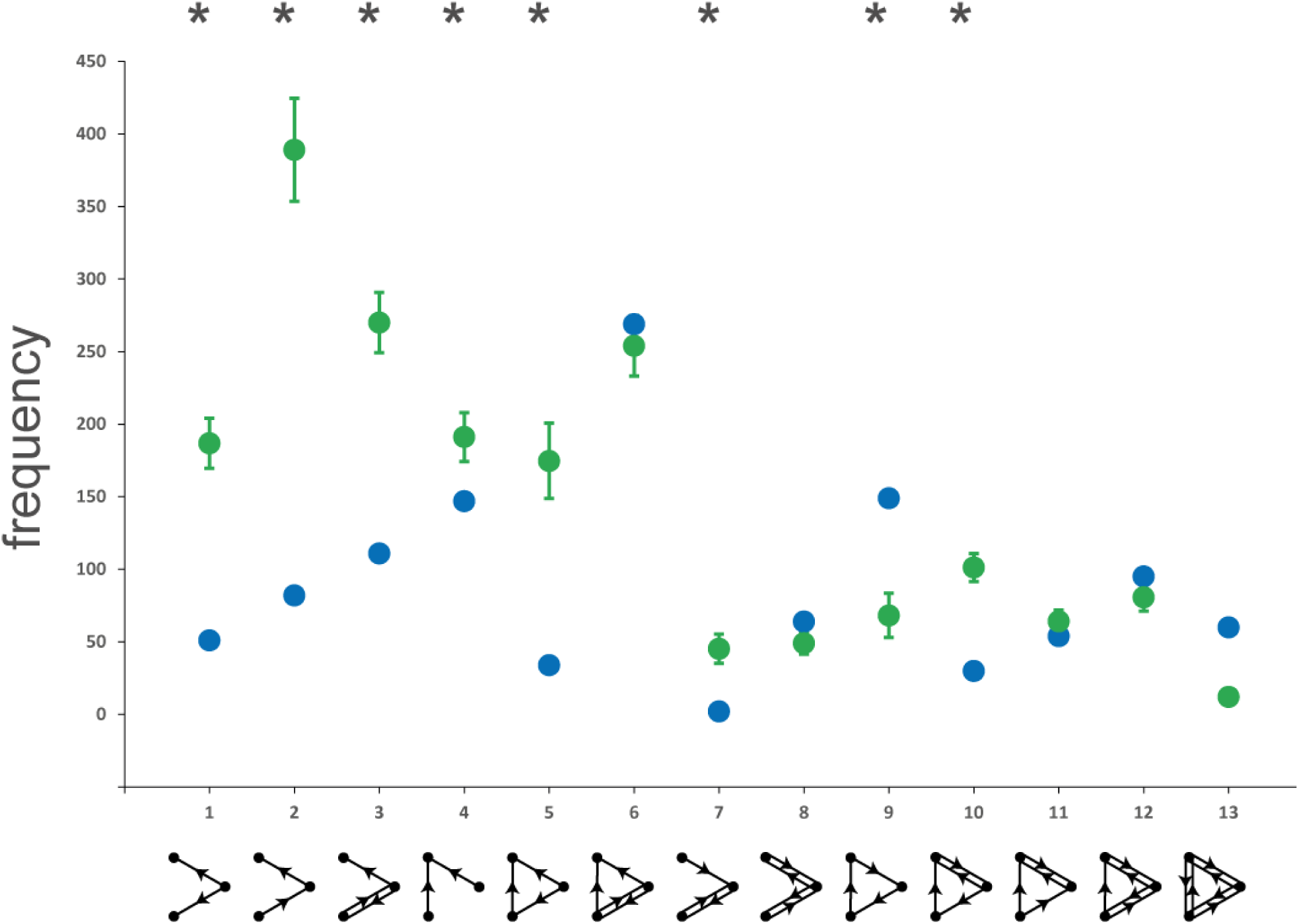
Motif composition and the 13 possible three-node motifs arrangements shown in the brain network of *O. vulgaris*. Network and mean (± SD) network motif fingerprint of the 1000 random networks of identical degree distributions are also shown. (B) Frequencies of the 13 possible 3-node motifs in the octopus’ network and mean (and SD) frequencies of the 1000 random networks. Asterisks denote significantly different frequencies (p < 0.05). (C) All 13 types of three-node patterns of connections (motifs); numbers refer to the ID. A single structural motif incorporates several functional motifs, for example, a structural motif (M = 3, ID = 13) may contain two separate instances of the motif ID = 9, or two motif ID = 7, or three distinct motif ID = 2.

The identification of motif spectrum is novel for any available study on cephalopod brains and supports the existence of a unique motif fingerprint, with specific non-random connection patterns. In *O. vulgaris* brain network, several motifs (9 and 13) exhibit a connectivity pattern in which all three nodes are bidirectionally interconnected, thereby supporting high levels of integration. This is observed for the optic lobe as well as the subvertical, medial basal, and lateral inferior frontal lobes, which also display properties of network hubs. It is noteworthy to underline the relative abundance of motifs featuring chains of reciprocally connected regions - without direct connections between their endpoints - suggesting a functional tendency to balance segregation and integration. This pattern may be associated with a specific type of neural dynamics (Tononi & Edelman, 1998; Tononi et al., 1998; Zhigulin, 2004).

## Discussion

This work characterizes the macroscale connectome of *Octopus vulgaris* brain using data on neural pathways and connectivity as described by Young (1971). We demonstrate that the octopus brain possesses multiple nonrandom attributes, including multiscale community structure, small-world properties, hubs, a unique motif fingerprint. These organizational principles are in line with numerous studies on other organisms, highlighting conserved architectural features across phylogeny, despite the phylogenetic distance between octopus and vertebrates.

The neural network analysis of *Octopus vulgaris* provides insights into the complexity and efficiency of the brain of this invertebrate animal. The study identified 439 neural connections, which were arranged in a binary matrix and represented afferent and efferent connections between brain lobes. Key hubs, including the medial basal lobe (32 links), subvertical lobe (31 links), and optic lobe (30 links), were identified by calculating the indegree and outdegree for each lobe and functional set. These regions play an essential role in the network, evidenced by the highest connectivity levels they exhibit.

The brain network of *O. vulgaris* diverges from random or scale-free networks. This topology has a small-world architecture with high clustering (0.64) and short path lengths (2.12) for effective information processing with low wiring cost. Furthermore, the network achieves great global and local efficiency by optimizing both the integration and segregation of information pathways.

The identification of community structures and modularity at various resolutions, is consistent with previously known functional groupings in the octopus brain, indicating both morphological and functional interconnectedness across brain areas. Notably, the optic lobe, basal lobes, and related areas form coherent clusters that demonstrate functional interdependence. A rich club arrangement was also identified, with high-degree hubs including the optic, subvertical, medial basal, prebrachial, and postbrachial lobes. These lobes exhibited highly dense interconnections, suggesting that they serve as a foundation for global communication and information integration across the brain.

Strikingly, the motif fingerprint of the octopus brain differs significantly from that of other invertebrates, such as *Caenorhabditis elegans*, and instead aligns more closely with mammalian neural networks (see Table 2). This resemblance highlights *O. vulgaris* brain remarkable capability to integrate and segregate information, which are essential for efficient and dynamic brain dynamics, as observed in complex neural networks.

**Table 2.** A comparative overview of small world features of the brain of O. vulgaris when compared with those known for other species. In the table: L= path length; C= clustering coefficient; N= number of nodes; d= connection density; λ= path-length scaled to random graph; γ= clustering coefficient scaled to random graph; σ=scalar small-worldness measure. Economic small-world parameters, global and local efficiency (Eglob and Eloc), and Cost (not computed for octopus). Note that for C. elegans, the number given corresponds to neurons and not areas. (*) number of neurons; (**) number of connections.

| Data | L | C | N | d | $\lambda$ | $\gamma$ | $\sigma$ | Eglob | Eloc | Cost |
| --- | --- | --- | --- | --- | --- | --- | --- | --- | --- | --- |
| *O. vulgaris* | 2.09 | 0.64 | 32 | 0.25 | 1.15 | 1.7 | 1.47 | 0.57 | 0.73 | - |
| *C. elegans* <sup>1</sup> | 2.65 | 0.28 | 282* |  | - | - | - | 0.45 | - | - |
| *D. melanogaster* <sup>2</sup> | 1.39 | 0.047 | 19,902* | 0.9 | - | 4.35 | 3.13 | - | - | - |
| *Human* <sup>3</sup> | 2.49 | 0.525 | 4005** | - | 1.09 | 2.37 | 2.18 | - | - | - |
| *Macaque* visual cortex <sup>4</sup> | 1.73 | 0.53 | 30 | - | 1.04 | 1.47 | - | - | - | - |
| *Macaque* whole cortex <sup>4</sup> | 2.38 | 0.46 | 39 | 0.15 | 1.17 | 3.06 | 2.44 | 0.52 | 0.7 | 0.18 |
| *Cat* cortex <sup>4</sup> | 1.81 | 0.55 | 65 | - | 1.06 | 1.77 | - | 0.69 | 0.83 | 0.38 |
| *Mouse* <sup>5</sup> | - | 0.159 | 111 | 53% | 1 | - | 1.31 | - | - | - |
| *Pigeon* telencephalon <sup>6</sup> | - | - | 52 | - | 2.40 | 0.36 | 1.04 | - | - | - |
| *Zebrafish* <sup>7</sup> | - | 0.132 | 419 | 2.45 | 2.86 | - | 4.13 | - | - | - |

1. Watts and Strogatz (1998)
2. Lin et al. (2024)
3. Achard et al. (2006)
4. Felleman and Van Essen (1991); Scannell et al. (1999); Young et al. (2000); Young et al. (2000); Sporns and Zwi (2004); Sporns et al. (2007)
5. Rubinov et al. (2015); Bassett and Bullmore (2016)
6. Shanahan et al. (2013)
7. Stobb et al. (2012)

Our results expand previous research that promoted the octopus brain as an ideal target organism for studying learning and memory, and the underlying neural machinery. Young (1964) suggested *O. vulgaris* brain as a "model of the models in the brain" and this has been further explored via the K.W. Craik’s proto-cybernetic theory of memory (Craik, 1967). These approaches were based on *O. vulgaris* predatory responses as originally characterized by Boycott, Maldonado, Packard and Young (Boycott, 1954; Boycott & Young, 1955; Maldonado, 1963a, 1963b; Packard, 1963). This cybernetic perspective has influenced several subsequent models of brain function inspired by the octopus. Marini et al. (2017) suggested that the cybernetic underpinnings of mid 20^th^ century memory models led to both broad initial adoption and eventual decline. For example, Clymer (1973) proposed a model in which visual features triggered memory values that guide a behavioral responses. Myers (1992) developed a neural network-based circuit designed to simulate learning responses of octopuses. These frameworks aimed to explain how information is processed and integrated to regulate complex actions in a brain with a relatively simple architecture (i.e., an invertebrate) yet capable of achieving sophisticated performance like that of mammals.

Recently, Chung et al. (2020; 2022) combined histological staining techniques and neural tracers with high resolution diffusion-weighted MRI to create a meso-scale atlas of the brain of *Sepioteuthis lessoniana* counting 426 connections between 23 different areas (Chung et al., 2020). Through a similar approach a connectivity map of the cuttlefish brain (*Sepia plangon*) was developed and compared to the ones of other cephalopod species known for possessing different lifestyles (*Vampyroteuthis infernalis*, *Hapalochlaena fasciata*, *Abdopus capricornicus* and *O. cyanea*, Chung et al., 2022).

Using high-resolution magnetic resonance imaging and fiber tractography, Jacobs (2022) mapped a mesoscale connectome of the *O. bimaculoides* brain, reporting segmentation of twenty-five lobes and tracing of fibers between them. The resulting connectivity matrix was largely consistent with findings from other investigations. Notably, due to the resolution used and the exceptionally small cell and fiber sizes, neural connectivity within the vertical lobe and its adjacent structures was not clearly evident from this study.

At the microscale level, Bidel et al. (2023) provided a synaptic-circuit analysis of the vertical lobe of *O. vulgaris*, thus greatly enriching our understanding of the structure of octopus neural networks.

By expanding previous studies (Gray & Young, 1964), Bidel and colleagues distinguished neuronal phenotypes: simple amacrine interneurons and more complex amacrine interneurons, with the former accounting for approximately 89.3% of the total cell population in the vertical lobe. Furthermore, unique connection patterns among interneurons have been identified, suggesting the presence of a complex architecture within the octopus vertical lobe.

The above mentioned studies illustrate lobe sizes, connections and information on neural cell types expanding previously comparative analysis of brains in this taxon (Nixon & Young, 2003; Ponte et al., 2021), and our knowledge on neural architecture of cephalopod brains. They also provide a further level of analysis, by enabling a future application of graph analysis (e.g., strength measures).

One of the key benefits of utilizing neural network analysis is its capacity to capture both global and local complexity. This approach enables the examination of interactions among different brain regions by taking into account both the overall (global) network structure and local dynamics. It also helps to identify critical nodes and hubs, i.e. brain areas that are both more central and/or influential (hubs) within the network and essential for overall brain function. Graph analysis enables the measurement of topological properties, providing metrics such as modularity, connectivity, and path length, which help to elucidate how the brain balances efficiency and robustness (Bullmore & Sporns, 2012).

Among invertebrates, the connectome of the nematode *Caenorhabditis elegans* provides an interesting case; with only 302 neurons and 2287 synaptic connections, *C. elegan*s was the first - and remains the most complete - invertebrate connectome mapped thus far (Varshney et al., 2011). By characterizing the entire wiring diagram of the animal, Authors contributed to exploring the relationship between structure and function of nervous systems (e.g., Chen et al., 2006). This seminal work sparked subsequent efforts to further map *C. elegans* nervous system and provided the basis for reconstructing local circuit connectomes of other worm species, i.e. *Platynereis dumerilii* (Jarrell et al., 2012; Randel et al., 2014).

After the first original work (Shih et al., 2015) *Drosophila* brain resulted at the center of an important effort in the characterization on the neural circuitry and connectomics (Lappalainen et al., 2024; Lin et al., 2024; Schlegel et al., 2024; Winding et al., 2023) for invertebrate brains.

Most of the studies on neural connectivity in vertebrates are based on mammalian species. Among various case studies is the mouse brain (see the Allen Mouse Brain Connectivity Atlas: https://connectivity.brain-map.org/ and: Oh et al., 2014; Wang et al., 2020; Zeng, 2018) applying various levels of accuracy from macro to microscale (e.g., Zhang et al., 2017), thus to provide important insights on how neural pathways contribute to sensory processing, motor control, and cognitive processes (e.g., Oh et al., 2014). Furthermore, the study of connectomes of the cerebral cortex in cats, monkeys, and humans has revealed shared topological features which have been linked to both normal and pathological behaviors (e.g., Bullmore & Bassett, 2011; Fornito et al., 2015; Goulas et al., 2014). These studies revealed how the cortical regions are connected with the surrounding areas in support of primary processing modules (e.g., visual, auditory, somato-motor, fronto-limbic), with a small number of direct, long-range pathways running between them. Among many features, small-world network organization has been detected, with highly connected regions involved in information processing, segregation and integration (e.g., Wang et al., 2021). Comparative connectomics has further advanced our understanding of neural networks by identifying conserved connectivity motifs and evolutionary changes in brain architecture. Collectively, these studies have enriched our understanding of how neural networks support behavior, cognition, and the development of neurological disorders (Barsotti et al., 2021; Galili et al., 2022; van den Heuvel et al., 2016).

While vertebrate connectomes provide insights into the complex functions and cognitive processes of the brain, invertebrate connectomes offer fundamental principles of neural connectivity that inform broader theories (Sporns, 2011), as well as valuable insights about the evolution of structural and functional neural complexity and broadly applicable principles of brain organization.

By gathering data from various species, connectomics continue to shed light on the complex relationship between brain structure and function across phylogeny.

The use of neural network analysis provides a more systemic perspective of nervous systems, moving beyond approaches that focus solely on specific connections or isolated regions. In addition, when combined with techniques such as MRI or HARDI, which allow for the assignment of weights to connections (for instance, based on the number of fibers), this type of analysis could help elucidate the importance of individual connections within the network and provide new insight on the relationship between different brain area.

In the present study, we applied neural network analysis to describe features of *O. vulgaris* brain. This is the first time such an approach has ever been used to characterize the structural properties of any cephalopod brain.

We identified characteristics (e.g., degree, motifs, rich-club) of the octopus network, and found analogous to those identified in mammalian brains. The identification of various brain regions appearing pivotal in the brain network such as the subvertical or medial basal lobes. These findings further supports the foundational knowledge acquired by Young, colleagues, and other researchers over decades of study.

Taken together, these results pave the way for an entirely novel approach to the analysis of cephalopod brain networks, as well as provide a foundation for future anatomical, functional, and behavioral studies in these organisms.

## Methods

All network metrics were computed on the basis of the octopus binary directed network using the Brain Connectivity Toolbox (Rubinov & Sporns, 2010), and additional sources as briefly illustrated below.

### Octopus brain binary connectivity

Data on macroscale white matter pathways of *O. vulgaris* brain was taken from Young (1971). Afferent and efferent connections reported between various areas and lobes of the octopus’ brain have been identified by J.Z. Young in the supraoesophageal (SEM), suboesophageal (SUB) masses and the optic lobes (OL). We did not consider nerves and other pathways departing from brain towards the periphery. Based on this information, a directed binary adjacent matrix of size N ⨯ N (N = 32) was constructed (Figure 1, Supplementary Info), expressing the presence of an anatomical projection between two regions with a 1 and absent and/or non-reported connections as a 0.

This matrix corresponded to a total of 32 nodes (see below for definitions), i.e. the cortical and subcortical regions within the octopus brain, and a collection of 2,178 hits reflecting the directed anatomical pathways between regions.

Aside from providing information on the existence or non-existence of neural pathways between regions in the *O. vulgaris* brain, our data did not permit the calculation of connectivity strength. Information on the number (and/or relative size) of fibers belonging to a given pathway or tract were not reported in the works we considered; available descriptions were limited to a few lobes but not consistently identified for all structures in the original studies (Budelmann & Young, 1985; Robertson et al., 1993; Young, 1971).

### Network metrics

We referred to Rubinov and Sporns (2010) for the graph analysis of the octopus connectivity matrix. We calculated network’s nodal degrees and degree distribution, i.e. number of afferent and efferent connections of brain regions. We considered the set of all n nodes in the network and all edges linking between nodes i and j. According to the original definition, links can be either undirected or directed. As mentioned previously and following the original description by J.Z. Young (1971), all links constituting the *O. vulgaris* brain network are directional. This allowed us to describe directionality: a connection projecting from a node (i) to another one (j) was considered bidirectional if projections from node j to node i exists; and unidirectional when no such connection was present.

We then calculated the degree distribution for all nodes, being the degree of node i the total number of connections belonging to it. In a similar fashion, in-degree of a node is equal to the total number of afferent connections of node i, and the out-degree corresponds to the total number of efferent connections of that node. As discussed by Rubinov and Sporns (2010) the degree of a given note reflects its relevance in a network. Bullmore and Sporns (2012) recommended that quantified network parameters should be compared with the null distribution of an equivalent random network. In random graphs each pair of nodes has an equal probability of being connected, reaching a Gaussian degree distribution in large networks. We assessed the degree distribution of *O. vulgaris* brain network against its equivalent random network (Rubinov & Sporns, 2010).

Modular organization of the octopus connectome was calculated using the algorithm (Newman’s modularity algorithm; Newman, 2006) available in the Brain Connectivity Toolbox (Rubinov & Sporns, 2010). This allows to estimate the modularity (Q) as whether nodes appearing in the same module are more often connected than in comparable randomly wired networks, with higher levels of Q indicating a more modular structure. We then linked the anatomical modules identified by the algorithm with specialized functional areas in the brain (e.g., chemo-tactile, visual regions). For the sake of this study, we assumed the existence of functional brain sets as originally described by Maddock and Young (1987). Authors grouped single brain lobes in eight functional sets and identified groupings of lobes (see Supplementary Info and Supplementary Table 5). For example: the vertical lobe system (VERT) includes the superior frontal, vertical, subvertical and precommissural lobes; the PALL counting on the palliovisceral and magnocellular lobes (Maddock & Young, 1987; sensu Young, 1971). Anatomical, functional, and effective modules in the octopus brain network show extensive overlap.

Participation coefficient between-modules is another measure informing about to what extent the connections of a given node are evenly distributed across the different modules of the network (Guimerà & Nunes Amaral, 2005), with nodes with a high participating coefficient reflecting regions with a high intermodular character. We measured assortativity: a measure of the tendency for nodes to be connected to other nodes of the same or similar degree. Positive assortativity indicates that high degree nodes tend to connect each other (see Box 2 in Bullmore & Sporns, 2012).

To capture the small-world organization of the *O. vulgaris* brain network, we assessed both clustering and characteristic path length. The clustering coefficient measures the likelihood that a node’s neighbours are interconnected by counting the fraction of triangles around each node. A low characteristic path length suggests that the network is well-configured for efficient signal transmission among constituent nodes. Also in this case, all measures were contrasted against a population of 1,000 randomly rewired networks with preserved in- and out-degree.

In the words of Bullmore and Sporns (2009) nearest neighbor nodes when connected to each other form a cluster. The clustering coefficient for a given node was calculated as the ratio of the actual number of connections between the neighbors of node 𝑖 to the total number of possible connections among those neighbors. The global clustering coefficient (𝐶) was then computed as the average of the clustering coefficients, 𝐶𝑖 for all nodes in the brain network.

Path length is the minimum number of edges that need to be traversed to travel from one node to another. In random and complex networks, the mean path length is short, indicating high global efficiency in parallel information transfer. Conversely, natural networks tend to have longer mean path lengths. Efficiency is inversely related to path length, but it is numerically easier to use for estimating topological distances between elements of disconnected graphs.

Path length is a purely topological concept and does not account for the physical length of a given connection. The characteristic (i.e. global) path length of the network is the average over path lengths for all nodes in the network (for definitions and discussion around small-world properties of a network see Rubinov & Sporns, 2010). The small-worldness organization of the network, was assessed considering clustering and characteristic path length.

Following Tononi (see: Tononi & Edelman, 1998; Tononi et al., 1994) anatomical brain connectivity is characterized by the simultaneous balance of the opposing of the opposing requirements for segregated/specialized and distributed/integrated information transfer and processing. An efficient anatomical network could thus include functionally specialized (segregated) modules as well as many intermodular (integrating) links. This type of architecture is typical of small-world networks and appears to be a common organization of anatomical neural connectivity (Bassett and Bullmore, 2006). Small-world networks are networks that are significantly more clustered than random networks while maintaining approximately the same characteristic path length (Watts & Strogatz, 1998).

Many small-world networks exhibit an economic property achieving high global and local efficiency while maintaining a low cost. This means that they effectively facilitate both long-distance communication and local clustering among nodes without requiring extensive resources. An estimator of the physical cost of a network is connection density, which is the ratio of the actual number of links to the total possible number of links (Bullmore & Sporns, 2009). Global efficiency is the inverse of characteristic path length and is more directly related to the system’s functional efficiency in parallel information transmission compared to characteristic path length (Bassett & Bullmore, 2006). The local efficiency is the global efficiency computed on the neighborhood of the node and is related to the clustering coefficient (Latora & Marchiori, 2001; Rubinov & Sporns, 2010).

The local efficiency of a network is a measure of the average efficiency of information transfer within local subgraphs (or neighborhoods) and is defined as the inverse of the shortest average path length of all neighbors of a given node among themselves. Local efficiency is an indication of how effectively information is integrated between the close neighbors of a given network node. On the other hand, global efficiency indicates how effectively information is integrated across the entirety of the network.

Small-world networks are formally defined as networks that are significantly more clustered than random networks while maintaining approximately the same characteristic path length. The shortest path length is the minimum number of edges traversed to get from a node to another node (Sporns et al., 2007).

To describe the balance of segregation and integration in the *O. vulgaris* brain network, we also investigated the presence of hubs in relation to their role in the community structure: provincial hubs are mainly connected to nodes in their own modules and are likely to promote modular segregation, while connector hubs are connected to nodes in other modules and are likely to facilitate global integration in the whole network For this purpose, we used the within-module degree z-score (z, a within-module version of degree centrality) and the participation coefficient (P), a measure of diversity of intermodular connections of individual nodes. The definitions of these two node categories were adopted from Guimerà and Nunes Amaral (2005; 2005).

A rich club is defined as a set of high-degree nodes that are also densely interconnected with each other, above and beyond their degree (Colizza et al., 2006; Opsahl et al., 2008). The procedure is performed over a range of degrees k. The rich club coefficient, R, at level k, is the fraction of links that connect nodes of degree k or higher out of the maximum number of links that such nodes might share. The normalized rich club coefficient (Rnorm) is the ratio between R, and the average rich club coefficient across random networks. The existence of a rich club organization is defined by Rnorm(k) > 1 over same range of values of threshold degree k. We utilized a probabilistic approach to define more precisely the threshold criteria for a rich club. At every different threshold degree, we estimated Rrandom(k) for 1000 different random networks and estimated the SD of Rrandom(k) (σ). The threshold range of the rich club regime was then specified by those values of k for which R(k) > Rrand(k) + 1σ. Therefore, a rich club could be said to exist in the subgroup of network nodes defined by an arbitrary degree threshold if Rnorm(k) > 1 + 1 σ.

Motifs are local connection patterns among a set of nodes and can be thought of as the building blocks of the network (Milo et al., 2002). As mentioned above, for three nodes there are 13 possible configurations, including cycles, open triangles, and so forth. Here we computed the frequency of each of the 13 three-node motifs, relative to their frequency in a population of 1,000 randomly rewired networks with preserved in- and out-degree sequence. A two-tailed p value (relative frequency of a particular motif) was estimated by computing the proportion of null networks with frequencies greater or smaller than in the real network.

We assessed bidirectionality by examining connections between nodes. A connection from node i to node j was classified as bidirectional if a corresponding projection from node j to node i was also present; otherwise, it was designated as unidirectional.

The betweenness centrality of a node i equals the normalized number of times node i is passed when walking along the shortest paths between all node pairs in the network, with nodes with the highest betweenness being among the most central nodes in the network.

## Acknowledgements

Federica Pizzulli has been supported by a PhD fellowship funded by the Stazione Zoologica Anton Dohrn (Open University – Stazione Zoologica Anton Dohrn PhD Program, XXII Cycle).

## Author contributions

F. Pizzulli: dataset compilation, contributed to formal analysis, writing of the original draft; M. Barjami: dataset compilation, formal analysis, contributed the application of a MatLab pipeline, and writing of the original draft; D. Edelman: writing, review and editing the original draft; G. Fiorito: conceptualization, writing, review and editing; G. Ponte: conceptualization, writing, review and editing. All Authors revised the final version of the manuscript.

