## Supplementary info for "Macroscale Connectivity in the Octopus brain"

#### Supplementary Information

Federica Pizzulli[1,#,*], Marisa Barjami[2,3,#], David B. Edelman[3], Graziano Fiorito[1], Giovanna Ponte[1,*]

[1] Department of Biology and Evolution of Marine Organisms, Stazione Zoologica Anton Dohrn, Villa Comunale, 80121 Napoli, Italy

[2] Dipartimento di Neuroscienze, Università di Parma, 43100 Parma, Italy

[3] Association for Cephalopod Research 'CephRes'-ETS, 80132 Naples, Italy

**Keywords**: Octopus; Brain Connectivity; Connectome; Motif; Hubs; Neural Network Analysis

[#] These Authors mutually contributed

[*] **Corresponding Authors**: Federica **Pizzulli**
Department of Biology and Evolution of Marine Organisms
Stazione Zoologica Anton Dohrn
Napoli, Italy


Giovanna **Ponte**
Department of Biology and Evolution of Marine Organisms
Stazione Zoologica Anton Dohrn
Napoli, Italy
Email: giovanna.ponte @szn.it

### The brain of *Octopus vulgaris* - a short overview

In the following pages, we first provide a short description of the overall organization of octopus nervous system.

The central brain of *O. vulgaris* consists of two masses (supra-esophageal mass, SEM; sub-esophageal mass, SUB) which are positioned above (SEM) and below (SUB) the anterior part of the oesophagus. A pair of optic lobes emerges from the sides of the supra-esophageal mass, behind the eyes (see below; Supplementary Figure 1). From the anterior region of the suboesophageal mass eight brachial nerves emerge, each one running in the center of an arm. In the arm, the axial nerve cord consists of two bundles of cerebro-brachial tracts (dorsal), and of a brachial ganglionic chain lying ventrally, made by an outer layer of cell bodies and of an internal neuropil.

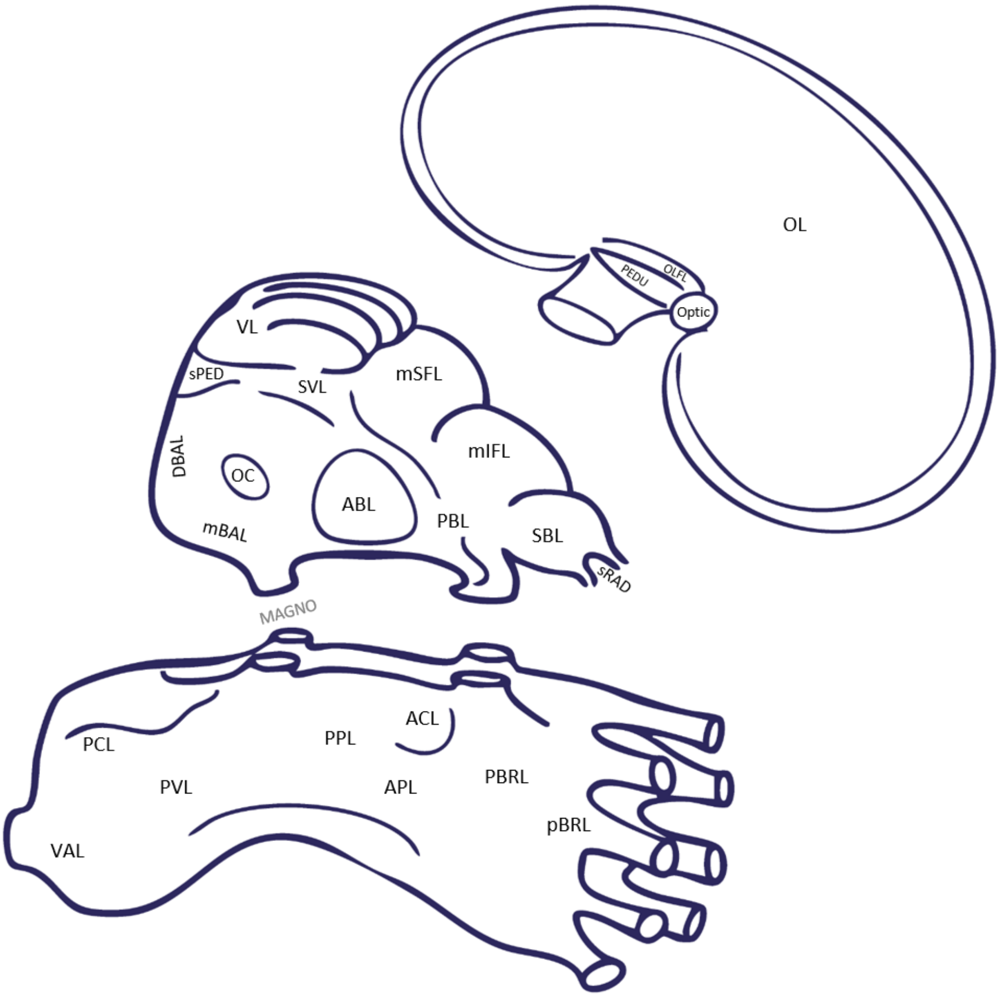

**Supplementary Figure 1**. An outline of *O. vulgaris* brain (supraoesophageal, suboesophageal and optic lobes). In the figure lobes are indicated and named (for list of abbreviations see Supplementary Table 1).

Each ganglion faces its corresponding sucker, thus appearing on alternate side of the cord with its typical sinusoidal arrangement (Graziadei, 1971; Margheri et al., 2011). The axial nerve cords (see also below) are linked to other peripheral nerve centers in the arm: **a.** four intramuscular nerve cords and **b.** the ganglia of the suckers. Sucker ganglia consist of a small cluster of cells, lying underneath the acetabular cup of each sucker (Graziadei, 1971). From the posterior end of the suboesophageal mass two pallial nerves emerge to connect the SUB to effectors in the mantle (i.e., the muscles and chromatophores) through the stellate ganglia.

A total of about 500 million nerve cells estimated to compose *O. vulgaris* nervous system; about 300 million are considered to belong to the arms' nervous components (Young, 1963). Supplementary Table 1 provides an overview of the distribution and numerosity of cells in various regions of the octopus brain.

**Supplementary Table 1.** Number of cells identified in the brain of *O. vulgaris* at various levels of distinct masses (Optic Lobes [OL], supra-oesophageal [SEM], sub-oesophageal [SUB]) and in the corresponding lobes/areas within each. Abbreviations of various lobes/parts in square brackets. Numbers in italics refer to structures included in a given lobe (see numbering as guidance).

| Structure | Lobe | cells x $10^3$ |
|---|---|---|
| **Optic** | | |
| | **Optic lobe [OL]** | **48000** |
| | **Peduncle lobe system** | |
| | Peduncle lobe(s) [PEDU] | 142 |
| | Olfactory lobe(s) [OLF] | 136 |
| | Optic gland(s) [Optic] | NA |
| **Supraoesophageal mass (SEM)** | | |
| | 1. Subradular ganglia [sRAD] | 32 |
| | 2. Superior buccal lobe [SBL] | 150 |
| | 3. Posterior buccal lobe [PBL] | na |
| | 4. Inferior buccal ganglia [IBL] | na |
| | 5. Inferior frontal lobe | 1085 |
| | 5.1. Lateral inferior frontal lobe [LatIFL] | *159* |
| | 5.2. Median inferior frontal lobe [mIFL] | *926* |
| | 6. Subfrontal lobe [SUBFL] | 5308 |
| | 7. Superior frontal lobe | 1854 |
| | 7.1. Lateral superior frontal lobe [SLFL] | *82* |
| | 7.2. Median superior frontal lobe [mSFL] | *1772* |
| | 8. Vertical lobe [VL] | 25066 |
| | 9. Subvertical lobe [SVL] | 810 |

| Structure | Lobe | cells x 10³ |
|---|---|---|
| | 10. Precommissural lobe [OC] | 78 |
| | 11. Anterior basal lobe [ABL] | 380 |
| | 12. Interbasal lobe [IntBAL] | 57 |
| | 13. Median basal lobe [mBAL] | 140 |
| | 14. Lateral basal lobe [LatBAL] | 127 |
| | 15. Dorsal basal lobe [DBAL] | 105 |
| | 16. Subpedunculate lobe [sPED] | 144 |
| **Suboesophageal mass (SUB)** | | |
| **Anterior Suboesophageal** | | 341 |
| | 1. Prebrachial lobe [pBRL] | 261 |
| | 2. Postbrachial lobe [PBRL] | 80 |
| **Middle Suboesophageal** | | 507 |
| | 1. Anterior pedal [APL], median | 135 |
| | 2. Lateral pedal [LatPL], | 36 |
| | 3. Posterior pedal [PPL], median | 72 |
| | 4. Anterior Chromatophore lobe [ACL] | 217 |
| | 5. Anterior funnel | 47 |
| **Posterior Suboesophageal** | | 2305 |
| | 1. Palliovisceral lobe [PVL] | 108 |
| | 2. Posterior chromatophore lobe [PCL] | 309 |
| | 3. Posterior funnel lobe | 47* |
| | 4. Vasomotor lobe [VAL] | 1307 |
| | 5. Magnocellular lobe [MAGNO] | 581 |

*anterio+ posterior funnel

The ‘central brain’ of *O. vulgaris* is included in a cartilaginous cranium (a sort of skull) that acts as a protection for the nervous tissue. The internal space created by this cranium is almost quite completely occupied by the brain, with the exception of the supra-esophageal mass that is projecting in the dorsal part of the cranium leaving a ‘surrounding’ space (about 60%) filled by a translucent jelly-like matrix. At the sonographic examination this space appears black (i.e. hypoechoic; Grimaldi et al., 2007).

The **supraoesophageal mass** (SEM) is composed by numerous lobes.

The ***superior buccal lobe***, the most anterior part of the SEM, contains 150 x 10³ cells (5–10 µm), with some exceeding 10 µm, reaching 20 µm (Young, 1963). Its posterior region connects to the ***inferior frontal lobe*** (1085 x 10³ cells), accounting for chemo-tactile information processing (i.e. memory systems/matrices, see: Young, 1991). The lobe is classically divided into lateral inferior frontal, median inferior frontal and sub-frontal lobes. These three parts are very dissimilar in terms of neural architecture. The medial part of inferior frontal lobe has a large neuropil and a thin layer of small cells (< 5 µm; with only a few much larger cells: 5 – 20 µm). The lateral inferior frontal lobe has many small cells (< 5 µm) and some large neurons (up to 20 µm). The ***sub-frontal lobe*** is principally composed by very small cells (nuclear diameter less than 3 µm) forming dense masses; dispersed in the wall of this lobe are cells with nuclei up to 20 µm.

The ***superior frontal lobe*** (1772 x $10^3$ cells) two regions are distinguished: lateral and medial superior frontal lobe. The first is composed by about half of cells < 5 µm (the remaining being 5 – 10 µm); the median superior frontal lobe is mainly composed by small cells (~ 4 µm).

The ***vertical lobe*** is the most dorsal and elongated structure of the supra-oesophageal mass. In *O. vulgaris* it is organized into five gyri (or lobules), i.e. approximately cylindrical structures running along the antero-posterior axis of the structure. Each 'cylinder' has a cellular wall surrounding a neuropil where cellular bodies are dispersed. The lobe is composed by a total of about 25,000 x $10^3$ cells; the larger number of cells is represented by small ones: amacrine, the smallest of octopus brain (3 µm in diameter). A much smaller number of larger cells with nuclear diameter 5 – 10 µm are also found. In contrast, with the usual arrangement of cells in the other lobes, the larger cells are closer to the neuropil.

Lying below the vertical lobe is a structure (**sub-vertical lobe**, 810 x $10^3$ cells) extending between the vertical, superior frontal, the basal lobes (ventral). It is characterized by the presence of a wall that in several regions is folded to form islands of cells (many < 5 µm, very few > 10 µm). At both sides of the sub-vertical lobe there are cells with diameter of 5 – 10 µm, but also neurons that are the largest of the whole supraesophageal mass (25 µm). In the center of the supraoesophageal mass and below the subvertical lobe, a structure made by many fibers is identified as the ***pre-commissural lobe*** (78 x $10^3$ cells), considered as a "meeting-point" for many fibre systems. It is characterized by few layers of cells, mostly of small ones (< 5 µm), and similar proportion of medium sized (5 – 10 µm) and some larger (around 1000 cells, 10 – 15 µm); its neuropil is continuous with the above lying sub-vertical lobe. More in a ventral position of the supra- oesophageal mass, are located a series of lobes belonging to the ***basal lobe system***. The basal 'system' is characterized by several structures (six lobes or parts) with cells of different dimensions and either distributed in layers or dispersed in the neuropile. Following Young (1971) classic works distinguished between 'anterior basal' and 'posterior basal' lobes. While the former is an easily recognizable entity, the 'posterior basal lobe' (i.e. 'occipital lobe' *sensu* Thore 1939; cited in Young, 1971) is a complex structure including other areas/lobes. Young adopts the term posterior basal lobe only in “a general sense for the large mass including all the above lobes except the anterior basal lobe” (Young, 1971, p. 349).

We distinguish (Young, 1963; Young, 1971):

***Anterior basal*** (380 x $10^3$ cells) with small or medium sized cells distributed in the dorsal and medial region of the lobe; at the level of the ventral region small, medium or larger cells are found (see also table 12.1 in Young, 1971). The lobe can be divided into three distinct subdivisions based on structure and connections: ***a.*** the “main part”, comprising the cell walls and neuropil found in the anterior region, as well as the posterior region’s hind part and lateral sides; ***b.*** a dorsal zone characterized by small cells; and ***c.*** a pair of antero-medial lobules containing notably large cells.

***Dorsal basal*** (2,000 x $10^3$ cells): with small and medium sized cells located at the level of the anterior part of the structure; cells of various sizes (< 5 µm, 5 – 10 µm, 10 – 15 µm) are found in the posterior region of the dorsal basal lobe. The dorsal basal lobe occupies the upper posterior portion of the supraoesophageal mass, below the vertical and rear regions of the subvertical lobes and above the median basal lobe. Many large cells have a main trunk extending downward, with few dendritic collaterals, likely contributing to efferent pathways to the optic lobes. Some trunks, however, turn upward, looping around cell islands. Small cells are often bipolar or multipolar, a characteristic shared by many large cells, resembling the optic lobe’s cellular organization. Emphasis is here provided also following Shigeno et al. (2018) who suggested that dorsal basal (and sub-vertical) as candidates for analogs to the vertebrate thalamus. This is based on many input fibers from the entire body via direct and indirect pathways from the sub-esophageal mass, suggesting it acts as relay center for the ‘cortically located’ other structures in cephalopod brain. Furthermore and according to Young (1971), the dorsal basal lobe contains large glial cells and lymphatic channels. These cells appear spherical or oval-shaped with spinous processes, while their tracts (in the dorsal region of the lobe) form bundles of long, fine fibres. Some originate in the posterior subvertical lobe and extend downward, while others loop from front to back, following axonal pathways. Certain fibres of these glia cells exceed 1 mm in length, making it difficult to differentiate them from nerve fibres, though tracing them to glial cell bodies confirms their attachment to a broad base with spinous processes.

***Subpedunculates*** (130 x $10^3$ cells; < 5 µm, 5 – 10 µm cells): this is one of the few supraoesophageal regions, apart from the superior buccal lobe, that sends nerve fibres outward the brain. It consists of two connected lobes, loosely distinguished from the dorsal basal lobe, to which it belongs as a specialized part. Positioned between the subvertical and vertical lobes, it contains multiple interconnected lobules of neuropil.

***Median basal*** (245 x $10^3$ cells of various sizes). The posterior basal part of the supraoesophageal lobes consists of a large neuropil mass with a thin outer nerve cell layer. The median basal lobe, centrally positioned above the oesophagus, maintains neuropil continuity with neighbouring lobes, suggesting functional connections. Despite this, its boundaries remain distinct. It lies beneath the dorsal basal lobes, with its dorsal limit marked by the ventral edge of the subvertical and dorsal basal lobes. Anteriorly, it connects broadly with the precommissural lobe, both above and below the ventral optic commissure. Ventrally, its neuropil extends to the interbasal, anterior basal, pedal, and palliovisceral lobes. Laterally and posteriorly, it merges with the lateral basal lobe (thus connecting towards chromatophore lobes, SUB). Above, the magnocellular lobes integrate with its sides (see also figure 14.5 in Young, 1971 for the connections of the lobe). A dorsal and ventral region are distinguished by Young.

***Interbasal*** (57 x $10^3$ cells of various sizes). It is a spherical structure, the most lateral and ventral part of the basal lobes, forming a distinct entity with unique cell structures and connections. Though its medial boundaries are difficult to define due to neuropil continuity with adjacent lobes, it remains clearly separate laterally. Positioned slightly above the oesophagus, it lies between the anterior basal, precommissural, and median basal lobes. Its lateral wall contains characteristic sets of large and small cells, while its neuropil forms a tangled network of fibres, some in loosely organized bundles. The interbasal lobes of both sides are likely connected by commissural fibres running ventral to the main optic commissure.

***Lateral basal*** (127 x $10^3$ cells of various sizes). The lateral basal lobes are considered by Young key components of the neural control-system of muscles of the chromatophores and skin. They are broadly continuous with the median basal but have distinct connections and characteristic cell layers and neuropil. It can be identified by a as slight swellings on the lower posterIor part of the supraoesophageal mass on either side of the median basal, with neuropil broadly continuous.

Outside the cranium and of each side of the head are the largest lobes of the whole central nervous system, i.e. the **optic lobes**. They contain about 92,000 x $10^3$ cells and represent about the 45% of the total volume of an octopus brain (e.g., Ponte et al., 2021). The optic lobes are characterized by the presence of an outer cortex and a central medulla. The cortex is composed by outer and inner layers of granular cells separated by a plexiform layer (i.e. neuropil). These layers contain cells smaller than 5 µm and several other medium sized cells (5 – 10 µm). The smaller cells are located next to the neuropil, while the larger ones lie at larger distance away from it. In the centre of the lobe, islands of nerve cells and many scattered small cells are found. Description of the ultrastructure and connections within the optic lobes is available from Saidel (1979; 1982). Proximal to the optic lobes and positioned on the optic tracts, the ***peduncle*** (142 x $10^3$ cells) and ***olfactory*** lobes (136 x $10^3$ cells) and the ***optic glands*** are found. The ***peduncle lobe*** is considered to be involved in the control of attack and movements (Messenger, 1967) and contains principally medium-sized cells (5 – 10 µm) with some others smaller than 5 µm. The ***olfactory lobe*** has a cellular composition like the peduncle lobe and seems to play a chemoreceptor function (the ***optic gland*** with endocrine function).

In the **subesophageal mass** (SUB) it is possible to distinguish three regions: **anterior**, **middle** and **posterior**. The anterior region is constituted by the **prebrachial** (261 x $10^3$ cells) and postbrachial lobes (80 x $10^3$ cells), involved in the control of arm movements. The prebrachial lobe is composed by a high proportion of small cells (5 µm) and large ones (20 µm in diameter); in contrast, in the postbrachial lobe large cells (25 µm) are the main constituents. The middle SUB region includes the **anterior chromatophore lobes** (217 x $10^3$ cells) and the pedal lobe (243 x $10^3$ cells). The higher proportion of neurons here is medium-sized (5 – 15 µm) or large cells (15 – 20 µm), and few smaller ones (< 5 µm). In contrast, the lateral parts of middle subesophageal mass contain a greater number of small cells. The posterior part of the middle SUB has pear-shaped cells (mainly medium in size). In the posterior SUB region are identifiable: the **palliovisceral** lobe (108 x $10^3$ cells), the **posterior chromatophore** lobes (309 x $10^3$ cells), **vasomotor** (1307 x $10^3$ cells) and **magnocellular lobes** (581 x $10^3$ cells). According to Young (1971), the palliovisceral lobe contains neurons that play different roles, but they are not separated in anatomically distinct districts. The outer cellular layer is composed by large cells (10 – 20 µm), instead near the neuropil there are very few small cells (5 µm). The vasomotor lobe has thick walls of numerous small cells (5 – 10 µm) with few large ones towards the periphery. The chromatophore lobes (one for each side) contain very numerous cells; in the most anterior part mainly small cells (5 – 10 µm) are present; in contrast, the most posterior part has 15 – 20 µm cells. The magnocellular lobes surround the oesophagous (one for each side) are considered to control the basic behavioural reactions including defensive motor patterns. The most dorsal part has small and medium-sized cells (5 - 10 µm) with only few larger cells (10 – 15 µm). In the ventral part the cell layers are much thicker and there are many larger cells (between 15 and 20 µm).

The **peripheral nervous system** of octopus is a complex network of nerves that, apart the nerve cord in the arms, has been described in great detail by Pfefferkorn (early XX century). Here we will not provide a throughout description of the peripheral nervous system of octopus; a detailed morphological overview is available in (Bullock, 1965) that also lists the nerves of the brain, their origin and relative position or role in the 'peripheral' network. It is useful to mention that Bullock describes ten series of nerves originating from the supraoesophageal mass and more than twenty from the suboesophageal mass (see table 25.1 in Bullock, 1965).
In addition, to the eight nerve cords in the arms and the ganglia at their bases (and the smaller ganglia occurring along the brachial nerves in the suckers), the peripheral system is composed by the following ganglia: the two stellate (one for each side of the mantle); two brachial, two or four (depending on the cephalopod species) cardiac. They are considered new developments in the cephalopods (Bullock, 1965). In each arm the nervous components can be summarized as follows:

i. a large axial nerve cord that runs through the centre of each arm;
ii. four thin nerve cords running longitudinally, near the periphery, in the
iii. muscular bundles of the arms;
iv. a sub-acetabular ganglion placed just in proximity of each sucker;
v. regular nerves emerging from each of above mentioned structures;
vi. small 'clumps' of nerve cells occur in the muscle;
vii. a notable number of nerve cells (probably sensory) superficially.

As mentioned, the arms of the octopus contain about two-thirds of the about 500 million neurons constituting the entire nervous system of the animal.

From the functional point of view, the nervous system of an octopus is characterized by a distributed organization due to the neural anatomy and elaborated sensory motor processing of the 'peripheral' system. The latter being wired to a 'central' system where coordination and decision-making units are present. The analysis of hundreds of lesion experiments conducted mainly on octopuses (review in Sanders, 1975) and of several dozens of serial histological sections of the brains of the animals allowed Young and coworkers to describe the functional anatomy of the nervous system by identifying a 'circuitry' leading to their visual and tactile processing: a circuit where learning and memory is achieved by a series of intersecting matrices (Young, 1991). An overview about these aspects is out of the aim of this work (see e.g., Marini et al., 2017; Ponte et al., 2022; Sanders, 1975; Young, 1991, 1995).

##### Neural connectivity in the brain of *O. vulgaris* – the dataset

Data on macroscale neural pathways of the octopus brain originated from the analysis and text mining of "The anatomy of the nervous system of *Octopus vulgaris*" by Young (1971).

These have been then transcribed and cross-tabulated (Supplementary Table 2) and a matrix with 1056 entries matching all reported connections within and/or between brain areas was compiled (Supplementary Figure 2). In the matrix connections were coded as presence (1) or absence (0) for the identification of a given tract/connection.

The resulting matrix represent the macroscale white matter pathways of the octopus brain.

| | OL | PEDU | OLFL | Optic | sRAD | IBL | SBL | PBL | LatIFL | mIFL | SUBFL | SLFL | mSFL | VL | SVL | ABL | OC | DBAL | sPED | mBAL | IntBAL | LatBAL | MAGNO | pBRL | PBRL | ACL | APL | LatPL | PPL | PVL | PCL | VAL |
|---|---|---|---|---|---|---|---|---|---|---|---|---|---|---|---|---|---|---|---|---|---|---|---|---|---|---|---|---|---|---|---|---|
| **OL** | 1 | 1 | 1 | | | | | | 1 | | | 1 | 1 | | 1 | 1 | 1 | 1 | 1 | 1 | 1 | 1 | 1 | | | | | 1 | 1 | | | |
| **PEDU** | 1 | 1 | 1 | | | | | | | | | | | | | 1 | | 1 | 1 | 1 | 1 | 1 | 1 | | | | | 1 | 1 | | | |
| **OLFL** | | 1 | 1 | 1 | | | | | | | | | | | | | | 1 | 1 | 1 | | 1 | | | | | | | | | | |
| **Optic** | | | | | | | | | | | | 1 | 1 | | | | | | | | | | | | | | | | | | | |
| **sRAD** | | | | | | | 1 | 1 | | | | | | | | | | | | | | | | | | | | | | | | |
| **IBL** | | | | | | | | | | | | | | | | | | | | | | | | | | | | | | | | |
| **SBL** | | | | | 1 | 1 | | 1 | 1 | 1 | 1 | | | | 1 | | | | | | | | | 1 | 1 | | | | | | | |
| **PBL** | | | | | | | 1 | | 1 | | 1 | | | | 1 | | | | | | | | | 1 | 1 | | | | | | | |
| **LatIFL** | 1 | | | | | | 1 | 1 | 1 | | | 1 | 1 | | 1 | | 1 | | | | | | | 1 | 1 | 1 | | | | | | |
| **mIFL** | 1 | | | | | | | | 1 | | 1 | | | | | | | | | | | | | | | | | | | | | |
| **SUBFL** | | | | | | | 1 | 1 | | 1 | | | | | | | | | | | | | | 1 | 1 | | | | | | | |
| **SLFL** | 1 | | | | | | | | 1 | | | 1 | 1 | 1 | 1 | | | | | | | | | | | | | | | | | |
| **mSFL** | 1 | | | | | | | | 1 | | | 1 | 1 | 1 | 1 | | | | | | | | | | | | | | | | | |
| **VL** | | | | | | | | | 1 | 1 | | 1 | 1 | 1 | 1 | | | | | | | | | | | | | | | | | |
| **SVL** | 1 | | | | | | 1 | 1 | 1 | 1 | 1 | 1 | 1 | 1 | 1 | | 1 | 1 | | 1 | | | 1 | | | | | | | | | |
| **ABL** | 1 | 1 | | | | | | | | | | | | | 1 | 1 | 1 | | | 1 | 1 | | | | | 1 | | 1 | 1 | | | |
| **OC** | | | | | | | 1 | | | | | | | | 1 | 1 | | | | 1 | 1 | | | | | 1 | | 1 | 1 | | | |
| **DBAL** | 1 | 1 | 1 | 1 | | | | | | | | | | | 1 | | | | 1 | 1 | | | 1 | | | | | | | | | |
| **sPED** | 1 | | | 1 | | | | | | | | | | | 1 | | | 1 | 1 | | | | | | | | | | | | | |
| **mBAL** | 1 | | | | | | | | | | | | | | 1 | 1 | 1 | 1 | 1 | 1 | 1 | 1 | 1 | | | | | 1 | 1 | 1 | 1 | 1 |
| **IntBAL** | | | | | | | | | | | | | | | | 1 | 1 | | | 1 | | | | | | | | 1 | | | | |
| **LatBAL** | 1 | | | | | | | | | | | | | | | 1 | | | | 1 | | | 1 | | | 1 | 1 | 1 | 1 | | 1 | |
| **MAGNO** | 1 | | | | | | | | | | | | | | 1 | | | | | 1 | | 1 | | 1 | 1 | | | 1 | 1 | 1 | | |
| **pBRL** | | | | | | | 1 | 1 | 1 | 1 | | 1 | 1 | 1 | 1 | 1 | 1 | | | 1 | 1 | | 1 | | 1 | 1 | | 1 | 1 | 1 | | 1 |
| **PBRL** | | | | | | | 1 | 1 | 1 | 1 | | 1 | 1 | 1 | 1 | 1 | 1 | | | 1 | 1 | | 1 | 1 | | 1 | | 1 | 1 | 1 | | 1 |
| **ACL** | | | | | | | | | | | | | | | | | | | | 1 | | | | 1 | 1 | | | 1 | 1 | | | |
| **APL** | | | | | | | | | | | | | | | | | | | | | | | | | | | 1 | | | | 1 | |
| **LatPL** | | | | | | | | | | | | | | | | | | | | 1 | | | 1 | | | 1 | | 1 | 1 | | | |
| **PPL** | | | | | | | | | | | | | | | | | | | | 1 | | | | | | 1 | | 1 | | 1 | | |
| **PVL** | | | | | | | | 1 | | | 1 | 1 | | 1 | 1 | 1 | | | | 1 | 1 | 1 | 1 | 1 | 1 | | | | 1 | | | 1 |
| **PCL** | | | | | | | | | | | | | | | | | | | | | | 1 | | | | | 1 | | | | 1 | |
| **VAL** | | | | | | | | | | | | | | | | | | | | | | | | 1 | 1 | | | | | 1 | | 1 |

**Supplementary Figure 2**. Presence/absence matrix depicting neural connections in various areas of *O. vulgaris* brain; afferent and efferent connections are reported after Young (1971). For abbreviations see Supplementary Table 1; See also Supplementary Table 2 and Figure 1, main text.

#### Supplementary Table 2 – list of tracts and connectives

**Supplementary Table 2**. List of tracts, connectives, commissures, nerves, and the entire network of fibres reported between lobes and areas of the *Octopus vulgaris* brain, as described by Young (1971). Afferent and efferent are distinguished by masses [Optic Lobe (OL), supraoesophageal (SEM) and suboesophageal (SUB) masses]. In the table we distinguish between neural connections reported to occur within lobes and masses, excluding those towards the periphery. The latter are summarized at the end of each table, but without listing the entire set of nerves emerging from the masses.

##### OPTIC LOBE (OL) – LIST OF NEURAL CONNECTIONS (WITHIN THE BRAIN)

| OL | Afferents | Efferents |
|---|---|---|
| **Optic Lobe** | Optic to **Optic lobes** | **Optic** to Optic lobes |
| | Peduncle to **Optic lobes** | **Optic** to Peduncle lobe |
| | Lateral inferior frontal to **Optic lobes** | **Optic** to Lateral inferior frontal lobes |
| | Median inferior frontal to **Optic lobes** | |
| | Lateral superior frontal to **Optic lobes** | **Optic** to Lateral superior frontal lobes |
| | Median superior frontal to **Optic lobes** | **Optic** to Median superior frontal lobes |
| | Subvertical to **Optic lobes** | **Optic** to Subvertical lobes |
| | Anterior basal to **Optic lobes** | **Optic** to Anterior basal lobes |
| | Dorsal basal to **Optic lobes** | **Optic** to Dorsal basal lobes |
| | Subpedunculate to **Optic lobes** | **Optic** to Subpedunculate lobes |
| | Median basal to **Optic lobes** | **Optic** to Median basal lobes |
| | Lateral basal to **Optic lobes** | **Optic** to Lateral basal lobes |
| | Magnocellular to **Optic lobes** | |
| | | **Optic** to Interbasal lobes |
| | | **Optic** to Lateral pedal lobes |
| | | **Optic** to Posterior pedal lobes |
| | | **Optic** to Precommissural lobes |
| **Peduncle lobe** | Peduncle to **Peduncle lobe** | **Peduncle** to Peduncle lobe |
| | Olfactory to **Peduncle lobe** | **Peduncle** to Olfactory lobe |
| | Anterior basal to **Peduncle lobe** | **Peduncle** to Anterior basal lobe |
| | Dorsal basal to **Peduncle lobe** | **Peduncle** to Dorsal basal lobe |
| | | **Peduncle** to Subpedunculate lobes |
| | | **Peduncle** to Median basal lobes |
| | | **Peduncle** to Interbasal lobes |
| | | **Peduncle** to Lateral basal lobes |
| | | **Peduncle** to Magnocellular lobes |
| | | **Peduncle** to Lateral pedal lobes |
| | | **Peduncle** to Posterior pedal lobes |
| | | **Peduncle** to Optic lobes |
| **Olfactory lobe** | Peduncle to **Olfactory lobe** | **Olfactory** to Peduncle lobes |
| | Olfactory to **Olfactory lobe** | **Olfactory** to Olfactory lobe |
| | Dorsal basal to **Olfactory lobe** | **Olfactory** to Dorsal basal lobes |
| | Optic to **Olfactory lobe** | **Olfactory** to Optic gland |

| OL | Afferents | Efferents |
|---|---|---|
| | | **Olfactory** to Subpedunculate lobes<br>**Olfactory** to Median basal lobes<br>**Olfactory** to Lateral basal lobes |
| **Optic gland** | Olfactory lobe to **Optic gland**<br>Dorsal basal to **Optic gland**<br>Subpedunculate to **Optic gland** | **Optic gland** to Lateral superior frontal lobes<br>**Optic gland** to Median superior frontal lobes |

**OPTIC LOBE (OL)** – LIST OF NEURAL CONNECTIONS TO OTHER MASSES (EXCLUDING SEM) AND EXTENDING TOWARD THE PERIPHERY

| Afferents | Efferents |
| --- | --- |
| Arms to Optic lobes | Optic lobes to Retina centrifugal fibres to the retina |
| Retina nerve fibres to Optic lobes | |

#### Supplementary Table 2 - *continued*

**SUPRAOESOPHAGEAL MASS (SEM)** – LIST OF NEURAL CONNECTIONS (WITHIN THE BRAIN)

| SEM - Lobes | Afferents | Efferents |
|---|---|---|
| **Subradular ganglia** | Superior buccal lobe to **Subradular ganglia** (cerebro-subradular connectives) | **Subradular ganglia** to Superior buccal lobe |
| | | **Subradular ganglia** to Posterior buccal lobe |
| **Superior Buccal lobes** | Subradular ganglia to **Superior buccal lobes** | **Superior buccal** to Subradular ganglia |
| | Posterior buccal to **Superior buccal lobes** | **Superior buccal** to Posterior buccal lobes |
| | Lateral inferior frontal to **Superior buccal lobes** | **Superior buccal** to Lateral inferior buccal tract |
| | Subfrontal to **Superior buccal lobes** | **Superior buccal** to Subfrontal lobes |
| | Subvertical to **Superior buccal lobes** | **Superior buccal** to Subvertical lobe |
| | Precommissural to **Superior buccal lobes** | |
| | Cerebrobrachial connective to **Superior buccal lobes** | **Superior buccal** to Brachial tract (Cerebrobrachial connective) |
| | | **Superior buccal** to Median inferior frontal lobes |
| | | **Superior buccal** to Inferior frontal lobes |
| **Posterior buccal lobes** | Inferior buccal to **Posterior buccal lobes** | |
| | Subradular ganglia to **Posterior buccal lobes** | |
| | Superior buccal to **Posterior buccal lobes** | **Posterior buccal** to Superior buccal lobes |
| | Subfrontal to **Posterior buccal lobes** | **Posterior buccal** to Subfrontal lobes |
| | Lateral inferior frontal to **Posterior buccal lobes** | **Posterior buccal** to Lateral inferior frontal lobes |
| | Subvertical to **Posterior buccal lobes** | **Posterior buccal** to Subvertical lobe |
| | Pre- and Postbrachial to **Posterior buccal lobes** | **Posterior buccal** to Pre- and Postbrachial lobes |
| | Pallioviscera to **Posterior buccal lobes** | |
| **Lateral inferior frontal lobes** | Optic to **Lateral inferior frontal lobes** | **Lateral inferior frontal** to Optic tract |
| | Superior buccal tract to **Lateral inferior frontal lobes** | **Lateral inferior frontal** to Superior buccal lobes |
| | Posterior buccal to **Lateral inferior frontal lobes** | **Lateral inferior frontal** to Posterior buccal lobes |
| | Lateral inferior frontal lobes to **Lateral inferior frontal lobes** | |
| | Median inferior frontal lobes to **Lateral inferior frontal lobes** | **Lateral inferior frontal** to Median inferior frontal lobes |
| | Lateral superior frontal lobes to **Lateral inferior frontal lobes** | **Lateral inferior frontal** to Lateral superior frontal lobes |

| SEM - Lobes | Afferents | Efferents |
| --- | --- | --- |
| | Median superior frontal lobes to **Lateral inferior frontal lobes** | **Lateral inferior frontal** to Median superior frontal lobes |
| | Vertical to **Lateral inferior frontal lobes** | |
| | Subvertical to **Lateral inferior frontal lobes** | **Lateral inferior frontal** to Subvertical lobes |
| | Pre- and Postbrachial to **Lateral inferior frontal lobes** (cerebro-brachial connective) | **Lateral inferior frontal** to Brachial lobes |
| | | **Lateral inferior frontal** to Precommissural lobes |
| | | **Lateral inferior frontal** to Anterior pedal lobes |
| **Median inferior frontal lobes** | Superior buccal to **Median inferior frontal lobes** | |
| | Subfrontal to **Median inferior frontal lobes** | **Median inferior frontal** to Subfrontal lobes |
| | Vertical to **Median inferior frontal lobes** | |
| | Subvertical to **Median inferior frontal lobes** | |
| | Pre- and Postbrachial to **Median inferior frontal lobes** (cerebro-brachial connective) | |
| | | **Median inferior frontal** to Optic Lobe |
| | | **Median inferior frontal** to Lateral inferior lobes |
| **Subfrontal lobes** | Superior buccal to **Subfrontal lobes** | **Subfrontal** to Superior buccal lobes |
| | Posterior buccal to **Subfrontal lobes** | **Subfrontal** to Posterior buccal lobes |
| | Median inferior frontal to **Subfrontal lobes** | **Subfrontal** to Median inferior frontal lobes |
| | Subvertical to **Subfrontal lobes** | |
| | | **Subfrontal** to Pre- and Postbrachial lobes |
| **Lateral superior frontal lobes** | Optic to **Lateral superior frontal lobes** | **Lateral superior frontal** to Optic lobe |
| | Optic gland to **Lateral superior frontal lobes** | |
| | Leteral inferior frontal to **Lateral superior frontal lobes** | **Lateral superior frontal** to Lateral inferior frontal lobes |
| | Lateral superior frontal to **Lateral superior frontal lobes** | **Lateral superior frontal** to Lateral superior frontal lobes |
| | Median superior frontal to **Lateral superior frontal lobes** | **Lateral superior frontal** to Median superior frontal lobes |
| | Vertical to **Lateral superior frontal lobes** | **Lateral superior frontal** to Vertical lobes |
| | Subvertical to **Lateral superior frontal lobes** | **Lateral superior frontal** to Subvertical lobes |
| | Pre- and Postbrachial to **Lateral superior frontal lobes** | |

| SEM - Lobes | Afferents | Efferents |
|---|---|---|
| | Palliovisceral to **Lateral superior frontal lobes** | |
| **Median superior frontal lobes** | Optic to **Median superior frontal lobes** | **Median superior frontal** to Optic lobe |
| | Optic gland to **Median superior frontal lobes** | |
| | Leteral inferior frontal to **Median superior frontal lobes** | **Median superior frontal** to Lateral inferior frontal lobes |
| | Lateral superior frontal to **Median superior frontal lobes** | **Median superior frontal** to Lateral superior frontal lobes |
| | Median superior frontal to **Median superior frontal lobes** | **Median superior frontal** to Median superior frontal lobes |
| | Vertical to **Median superior frontal lobes** | **Median superior frontal** to Vertical lobes |
| | Subvertical to **Median superior frontal lobes** | **Median superior frontal** to Subvertical lobes |
| | Pre- and Postbrachial to **Median superior frontal lobes** | |
| **Vertical lobes** | Lateral superior frontal to **Vertical lobes** | **Vertical** to Lateral superior frontal lobes |
| | Median superior frontal to **Vertical lobes** | **Vertical** to Median superior frontal lobes |
| | Vertical to **Vertical lobes** | **Vertical** to Vertical lobes |
| | Subvertical to **Vertical lobes** | **Vertical** to Subvertical lobes |
| | Pre- and Postbrachial to **Vertical lobes** | |
| | Palliovisceral to **Vertical lobes** | |
| | | **Vertical** to Lateral inferior frontal lobes |
| | | **Vertical** to Median inferior frontal lobes |
| **Subvertical lobes** | Optic to **Subvertical lobes** | **Subvertical** to Optic lobes |
| | Superior buccal to **Subvertical lobes** | **Subvertical** to Superior buccal lobes |
| | Posterior buccal to **Subvertical lobes** | **Subvertical** to Posterior buccal lobes |
| | Lateral inferior frontal to **Subvertical lobes** | **Subvertical** to Lateral inferior frontal lobes |
| | Lateral superior frontal to **Subvertical lobes** | **Subvertical** to Lateral superior frontal lobes |
| | Median superior frontal to **Subvertical lobes** | **Subvertical** to Median superior frontal lobes |
| | Vertical to **Subvertical lobes** | **Subvertical** to Vertical lobes |
| | Subvertical to **Subvertical lobes** | **Subvertical** to Subvertical lobes |
| | Anterior basal to **Subvertical lobes** | |
| | Precommissural to **Subvertical lobes** | **Subvertical** to Precommissural lobes |

| SEM - Lobes | Afferents | Efferents |
| --- | --- | --- |
| | Dorsal basal to **Subvertical lobes** | **Subvertical** to Dorsal basal lobes |
| | Subpedunculate to **Subvertical lobes** | |
| | Median basal to **Subvertical lobes** | **Subvertical** to Median basal lobes |
| | Magnocellular to **Subvertical lobes** | **Subvertical** to Magnocellular lobes |
| | Pre- and Postbrachial to **Subvertical lobes** | |
| | Palliovisceral to **Subvertical lobes** | |
| | | **Subvertical** to Median inferior frontal lobes |
| | | **Subvertical** to Subfrontal lobes |
| **Anterior basal lobes** | Optic to **Anterior basal lobes** | **Anterior basal** to Optic lobes |
| | Peduncle to **Anterior basal lobes** | **Anterior basal** to Peduncle lobes |
| | Anterior basal to **Anterior basal lobes** | **Anterior basal** to Anterior basal lobes |
| | Precommissural to **Anterior basal lobes** | **Anterior basal** to Precommissural lobes |
| | Median basal to **Anterior basal lobes** | **Anterior basal** to Median basal lobes |
| | Interbasal to **Anterior basal lobes** | **Anterior basal** to Interbasal lobes |
| | Lateral basal to **Anterior basal lobes** | **Anterior basal** to Lateral pedal lobes |
| | Pre- and Post brachial to to **Anterior basal lobes** | |
| | Palliovisceral to to **Anterior basal lobes** | |
| | | **Anterior basal** to Subvertical lobes |
| | | **Anterior basal** to Anterior pedal lobes |
| | | **Anterior basal** to Posterior pedal lobes |
| **Precommissural lobes** | Optic to **Precommissural lobes** | |
| | Lateral inferior frontal to **Precommissural lobes** | |
| | Subvertical to **Precommissural lobes** | **Precommissural** to Subvertical lobes |
| | Anterior basal to **Precommissural lobes** | **Precommissural** to Anterior basal lobes |
| | Median basal to **Precommissural lobes** | **Precommissural** to Median basal lobes |
| | Interbasal to **Precommissural lobes** | **Precommissural** to Interbasal lobes |
| | Pre- and Postbrachial to **Precommissural lobes** | |

| SEM - Lobes | Afferents | Efferents |
|---|---|---|
| | | **Precommissural** to Superior buccal lobes |
| | | **Precommissural** to Anterior pedal lobes |
| | | **Precommissural** to Lateral pedal lobes |
| | | **Precommissural** to Posterior pedal lobes |
| **Dorsal basal lobes** | Optic to **Dorsal basal lobes** | **Dorsal basal** to Optic lobes |
| | Peduncle to **Dorsal basal lobes** | **Dorsal basal** to Peduncle lobes |
| | Olfactory to **Dorsal basal lobes** | **Dorsal basal** to Olfactory lobes |
| | Subvertical to **Dorsal basal lobes** | **Dorsal basal** to Subvertical lobes |
| | Subpedunculate to **Dorsal basal lobes** | **Dorsal basal** to Subpedunculate lobes |
| | Median basal to **Dorsal basal lobes** | **Dorsal basal** to Median basal lobes |
| | | **Dorsal basal** to Optic glands |
| | | **Dorsal basal** to Magnocellular lobes |
| **Subpeduncualte lobes** | Optic to **Subpedunculate lobes** | **Subpedunculate** to Optic lobes |
| | Peduncle to **Subpedunculate lobes** | |
| | Olfactory to **Subpedunculate lobes** | |
| | Dorsal basal to **Subpedunculate lobes** | **Subpedunculate** to Dorsal basal lobes |
| | Subpedunculate to **Subpedunculate lobes** | **Subpedunculate** to Subpedunculate lobes |
| | Median basal to **Subpedunculate lobes** | |
| | | **Subpedunculate** to Optic glands |
| | | **Subpedunculate** to Subvertical lobes |
| **Median basal lobes** | Peduncle to **Median basal lobes** | |
| | Olfactory to **Median basal lobes** | |
| | Dorsal basal to **Median basal lobes** | **Median basal** to Dorsal basal lobes |
| | Subvertical to **Median basal lobes** | **Median basal** to Subvertical lobes |
| | Anterior basal to **Median basal lobes** | **Median basal** to Anterior basal lobes |
| | Precommissural to **Median basal lobes** | **Median basal** to Precommissural lobes |
| | Madian basal to **Median basal lobes** | **Median basal** to Median basal lobes |

| SEM - Lobes | Afferents | Efferents |
|---|---|---|
| | Interbasal to **Median basal lobes** | **Median basal** to Interbasal lobes |
| | Lateral basal to **Median basal lobes** | **Median basal** to Lateral basal lobes |
| | Magnocellular to **Median basal lobes** | **Median basal** to Magnocellular lobes |
| | Pre- and Postbrachial to **Median basal lobes** | |
| | Anterior pedal to **Median basal lobes** | |
| | Lateral pedal to **Median basal lobes** | **Median basal** to Lateral pedal lobes |
| | Posterior pedal to **Median basal lobes** | **Median basal** to Posterior pedal lobes |
| | Palliovisceral to **Median basal lobes** | **Median basal** to Palliovisceral lobes |
| | | **Median basal** to Optic lobes |
| | | **Median basal** to Subpedunculate lobes |
| | | **Median basal** to Vasomotor lobes |
| | | **Median basal** to Posterior chromatophore lobes |
| **Interbasal lobes** | Optic to **Interbasal lobes** | |
| | Peduncle to **Interbasal lobes** | |
| | Anterior basal to **Interbasal lobes** | **Interbasal** to Anterior basal lobes |
| | Precommissural to **Interbasal lobes** | **Interbasal** to Precommissural lobes |
| | Median basal to **Interbasal lobes** | **Interbasal** to Median basal lobes |
| | Pre- and Postbrachial to **Interbasal lobes** | |
| | Palliovisceral to **Interbasal lobes** | |
| | | **Interbasal** to Lateral pedal lobes |
| **Lateral basal lobes** | Optic to **Lateral basal lobes** | **Lateral basal** to Optic lobes |
| | Peduncle to **Lateral basal lobes** | |
| | Olfactory to **Lateral basal lobes** | |
| | Median basal to **Lateral basal lobes** | **Lateral basal** to Median basal lobes |
| | Magnocellular to **Lateral basal lobes** | **Lateral basal** to Magnocellular lobes |
| | Posterior pedal to **Lateral basal lobes** | **Lateral basal** to Posterior pedal lobes |
| | Palliovisceral to **Lateral basal lobes** | |
| | Posterior chromatophore lobes to **Lateral basal lobes** | **Lateral basal** to Posterior chromatophore lobes |
| | | **Lateral basal** to Anterior chromatophore lobes |

| SEM - Lobes | Afferents | Efferents |
|---|---|---|
| | | **Lateral basal** to Lateral pedal lobes<br>**Lateral basal** to Anterior basal lobes<br>**Lateral basal** to Anterior pedal lobes |

**SUPRAOESOPHAGEAL MASS (SEM)** – LIST OF NEURAL CONNECTIONS EXTENDING TOWARD THE PERIPHERY

| Afferents | Efferents |
|---|---|
| Lips to Superior buccal lobes | Superior buccal to Lips |
| Buccal mass to Superior buccal lobes | Superior buccal to Arms |
| Arms to Superior buccal lobes | Superior buccal to Posterior salivary glands (33000 fibres) |
| Posterior salivary glands to Superior buccal lobes | Superior buccal to Cerebro-subradular connective (6600 fibres) |
| Posterior salivary glands to Subradular ganglion | Superior buccal to Interbuccal connective |
| Radular nerves to Inferior buccal lobes | Inferior buccal to Superior mandibular nerves |
| Sympathetic nerves to Inferior buccal ganglia | Inferior buccal to Lateral buccal palps |
| Arms to inferior frontal lobes | Inferior buccal ganglia to Anterior salivary glands |
| | Inferior buccal ganglia to Sympathetic nerves |
| | Inferior buccal ganglia to Juxta-ganglionic tissue |

**Supplementary Table 2** *- continued*

**SUBOESOPHAGEAL MASS (SUB)** – LIST OF NEURAL CONNECTIONS (WITHIN THE BRAIN)

| SUB - Lobes | Afferents | E |
|---|---|---|
| **Anterior Pedal lobes** | Brachial to **Anterior pedal tract** | **Anterior pedal** to Brachia |
| | Anterior basal to **Anterior pedal tract** | |
| | Lateral pedal to **Anterior pedal tract** | |
| | Posterior pedal to **Anterior pedal tract** | **Anterior pedal** to Posteri |
| | Precommissural to **Anterior pedal tract** | **Anterior pedal** to Precom |
| | Lateral basal to **Anterior pedal tract** | |
| | Lateral inferior frontal to **Anterior pedal tract** | |
| | | **Anterior pedal** to Lateral |
| | | **Anterior pedal** to Precom |
| | | **Anterior pedal** to Antorb |
| | | **Anterior pedal** to Median |
| **Lateral pedal lobes** | Optic to **Lateral pedal lobes** | |
| | Peduncle to **Lateral pedal tract** | |
| | Anterior basal to **Lateral pedal tracts** | |
| | Median basal to **Lateral pedal tracts** | **Lateral pedal** to Median |
| | Precommissural to **Lateral pedal tract** | |
| | Basal lobes to **Lateral pedal lobes** | |
| | Magnocellular to **Lateral pedal tract** | **Lateral pedal** to Magnoce |
| | Bachial to **Lateral pedal tract** | |
| | Anterior pedal to **Lateral pedal tract** | **Lateral pedal** to Anterior |
| | Lateral pedal to **Lateral pedal tract** | **Lateral pedal** to Lateral p |
| | Posterior pedal to **Lateral pedal tract** | **Lateral pedal** to Posterior |
| **Posterior pedal lobes** | Optic to **Posterior pedal tract** | |
| | Peduncle to **Posterior pedal tract** | |
| | Anterior basal to **Posterior pedal lobes** | |
| | Precommissural to **Posterior pedal lobes** | |
| | Lateral basal lobes to **Posterior pedal tract** | |
| | Median basal lobes to **Posterior pedal tract** | **Posterior pedal** to Media |
| | Magnocellular to **Posterior pedal tract** | |
| | Anterior pedal to **Posterior pedal tract** | **Posterior pedal** to Anteri |
| | Bachial to **Posterior pedal tract** | |
| | Lateral pedal to **Posterior pedal tract** | **Posterior pedal** to Latera |
| | Palliovisceral to **Posterior pedal tract** | **Posterior pedal** to Palliov |
| **Anterior chromatophore lobes** | Lateral basal to **Anterior chromatophore lobes** | |
| | Posterior chromatophore to **Anterior chromatophore lobes** | **Anterior chromatophore** lobes |
| | Anterior chromatophore to **Anterior chromatophore lobes** | **Anterior chromatophore** lobes |
| | Posterior Suboesophageal | |
| | Includes: | 1. Palliovisceral lobes |
| | | 2. Postero-dorsal lobes |
| | | 3. Posterior chromatopho |
| | | 4. Posterior funnel lobes |

| SUB - Lobes | Afferents | E |
|---|---|---|
| | | 5. Vasomotor lobes |
| | | 6. Magnocellular lobes, fo |
| | **Afferent** | **Efferent** |
| **Palliovisceral tract** | Median Basal to **Palliovisceral tract** | **Palliovisceral** to Median I |
| | Magnocellular to **Palliovisceral tract** | **Palliovisceral** to Magnoce |
| | Brachial to **Palliovisceral tract** | **Palliovisceral** to Pre- and |
| | Posterior Pedal to **Palliovisceral tract** | **Palliovisceral** to Posterio |
| | Vasomotor to **Palliovisceral lobes** | **Palliovisceral** to Vasomot |
| | Pedal to **Palliovisceral lobes** | |
| | | **Palliovisceral** to Anterior |
| | | **Palliovisceral** to Subfront |
| | | **Palliovisceral** to Interbas |
| | | **Palliovisceral** to Lateral b |
| | | **Palliovisceral** to Superior |
| | | **Palliovisceral** to Vertical I |
| | | **Palliovisceral** to Subverti |
| | | **Palliovisceral** to Posterio |
| **Posterior chromatophore lobes** | Median Basal to **Posterior chromatophore lobes** | |
| | Lateral Basal to **Posterior chromatophore lobes** | **Posterior chromatophore** |
| | Anterior chromatophore to **Posterior chromatophore lobes** | **Posterior chromatophore** |
| | Posterior chromatophore to **Posterior chromatophore lobes** | **Posterior chromatophore** lobes |
| **Vasomotor tract** | Median Basal to **Vasomotor tract** | |
| | Brachio-palliovisceral tract to **Vasomotor lobes** | **Vasomotor** to Pre- and P |
| | Palliovisceral to **Vasomotor lobes** | **Vasomotor** to Palliovisce |
| | Vasomotor to **Vasomotor lobes** | **Vasomotor** to Vasomotor |
| **Magnocellular tract** | Optic to **Magnocellular tract** | **Magnocellular** to Optic |
| | Peduncle to **Magnocellular tract** | |
| | Subvertical to **Magnocellular tract** | **Magnocellular** to Subvert |
| | Dorsal basal to **Magnocellular tract** | |
| | Median basal to **Magnocellular tract** | **Magnocellular** to Median |
| | Lateral basal to **Magnocellular tract** | **Magnocellular** to Lateral |
| | Brachio-palliovisceral connective to **Magnocellular tract** | **Magnocellular** to Brachio |
| | Lateral pedal to **Magnocellular lobes** | **Magnocellular** to Lateral |
| | | **Magnocellular** to Posteri |
| | | **Magnocellular** to Palliovi |

**SUBOESOPHAGEAL MASS (SUB)** – LIST OF NEURAL CONNECTIONS EXTENDING TOWARD THE PERIPHERY

| Afferents | Efferents |
|---|---|
| Arms to Postbrachial lobes | Postbrachial to Arms |

Antorbital to Prebrachial lobes

Statocysts  to lateral pedal lobes
Pallial nerves to Palliovisceral lobes
Collar nerves to Palliovisceral lobes
Visceral nerves to Palliovisceral lobes
Heart and other viscera to Vasomotor lobes
Pallial nerve to Magnocellular tract
Anterior infundibular to Magnocellular lobes
Interbrachial nerves to Brachial lobes
Ophthalmic to lateral pedal nerves
Static nerve to middle pedal commissure
Static nerve to magnocellular lobe
Static nerve to palliovisceral lobe
Static nerve to median basal lobe
Static nerve to anterior basal lobe

Posterior basal-suboesophageal to Median basal lo
Pedal to Antorbital nerves (oculomotor and ophtha
nerves)
Pedal lobes to Arms
Pedal lobes to Anterior funnel nerves (800 fibres)
Pedal lobes to head retractor muscles
Posterior pedal to Median pallial retractor nerves
Palliovisceral lobes to Pallial nerves (5500 fibres)
Palliovisceral lobes to Collar nerves (1800 fibres)
Palliovisceral lobes to Funnel ( ~2000 fibres)
Palliovisceral lobes to Visceral (4400 fibres)
Vasomotor lobes to musculature of the arteries and
Vasomotor lobes to Anterior vena cava nerve (1000
Vasomotor lobes to Visceral nerves (52000 fibres)
Vasomotor lobes to Perioesophageal nerves (aortic

#### Estimation of fibres in the *O. vulgaris* brain

As mentioned, the number of directed links here accounted is underestimated: we did not include in this analysis paths or connections reaching the peripheral nervous system or targets external to the brain (e.g., ophthalmic and oculomotor nerves, mandibular- radular and salivary nerves, visceral and cardiac nerves; see also Supplementary Table 2).

Furthermore, the original dataset did include, only in a few instances, information on the number of fibres for a given tract or neural pathway in the neuropile (see Young, 1971). Thus, we avoided any estimation of edge strength to avoid potential bias.

As an example, about 16,000 fibres have been estimated in each octopus pallial nerve (Budelmann & Young, 1985); these are distinguished as follows:

a. around 3200 fibres (mostly of those with a diameter of 2-300 6 µm) make synapses with the motor neurons inside the stellate ganglion, which then innervate the respiratory mantle muscles;
b. 600 fibres (between 6-16 µm in diameter) directed to the head retractor muscles, leaving the nerve before reaching the stellate ganglion;
c. about 2300 fibres (less than 4 µm in diameter) run into the ganglion and exit through the stellate nerves to directly innervate chromatophores in the skin;
d. around 10000 extremely fine fibres (< 1 µm and/or < 2 µm in diameter) identified as afferents originating from the periphery.

Centripetal cells were also identified inside the stellate ganglion (Monsell, 1977; Monsell, 1980), with the axons pointing toward the pallial nerve. Their targets have never been identified and they have been considered sensory neurons, which may serve for transmitting information from the sub-stellar organs, the latter considered being stretch receptors. Supplementary Table 3 reports examples of the estimation of neural fibres, as originally included in the studies.

#### Supplementary Table 3 - fibers

Examples of number of fibres in various locations of the nervous system of *O. vulgaris* as accounted by Young and colleagues (Budelmann & Young, 1985; Young, 1965; Young, 1971).

| Areas/Lobes | Fibres |
|---|---|
| **Peduncle commissure** | 3000 |
| **Pallial nerve** | 5500 |
| **Brachial nerve:** lateral motor root | 6300 |
| **Brachial (one side)** | 72000 |
| other areas of the lateral root | 1600 |
| **Cerebro-brachial tract** | 30000 |
| **Pedal-Palliovisceral** | 280 |
| **Lateral pedal – Oculomotor** | 927 |
| **Posterior Pedal – Anterior funnel** | 800 |
| **Visceral – Vasomotor** | 50000 |
| **Cerebro - subradular connective** | 6600 |
| **Brain to Stellate ganglia** | 4000 |

### Functional Brain Sets in the octopus brain

Young and coworkers grouped single brain lobes in functional sets (L. Maddock & J. Z. Young, 1987). The original aim was to facilitate comparisons between brain organization of different cephalopod species. In the original set the vertical lobe system (VERT, Supplementary Table 5) includes the superior frontal, vertical, subvertical and precommissural lobes (L. Maddock & J.Z. Young, 1987; sensu Young, 1971). In order to facilitate an analysis of the neural network approach amd search for possible functional correlation with modularity and other parameters, we adopted the functional sets approach of (L. Maddock & J. Z. Young, 1987); a similar approach has been followed a comparative analysis of cephalopod brain organization in this taxon (Ponte et al., 2021).

#### Supplementary Table 4 – Functional sets of octopus brain

List of the eight functional sets and corresponding octopus brain lobes. The work of L. Maddock and J.Z. Young (1987) is here considered since the authors introduced the concept of “functional sets” to the taxon, to represent the various lobes of the brain in various species of cephalopods.

| Functional sets | Maddock and Young, 1987 |
|---|---|
| ***Supraoesophageal mass*** | |
| INFF | Inferior frontal<br>Subfrontal<br>Superior buccal<br>Median inferior frontal |
| VERT | Superior frontal<br>Vertical<br>Subvertical<br>Precommissural |
| BASAL (=PARA)[3] | Anterior basal<br>Peduncle<br>Olfactory |
| ***Supraoesophageal mass** - continued* | |
| BASAL (=MEDB)[3] | Median basal<br>Dorsal basal<br>Interbasal<br>Dorsolateral<br>Subpedunculate |
| BASAL (=LATB)[3] | Lateral basal<br>Optic gland[4] |
| ***Suboesophageal mass*** | |
| BRAC | Brachial |
| PEDAL | Pedal |
| PALL | Palliovisceral<br>Magnocellular |
| CHRF[5] | Chromatophore[6]<br>Fin |
| ***Optic lobes*** | |
| OPTIC | Optic |

4. The optic gland is listed in Table II of Maddock and Young (1987), but is not included in the equivalent list of Nixon and Young (2003).

5. Following Wirz (1959), the values of the chromatophore and fin lobes were summed up together in the same functional set (here called CHRF) and not considered separately as CHROM and FINL (*sensu:* Maddock and Young, 1987; Nixon and Young, 2003).

6. As reported at p. 758 of Maddock and Young (1987), it should be the posterior chromatophore lobe.

#### Degree

We calculated the degree of a node i as the total number of connections linked to it. Similarly, the in-degree of node i represents the total number of incoming connections it receives. As discussed in main text afferences and efferences in *O. vulgaris* brain are not mutually equivalent. In particular we found: eighteen octopus brain regions having more afferences than efferences, if considering areas classified as PER i.e. belonging to SEM: 11 nodes, SUB: 7 nodes, ten (10) nodes show the opposite trend (outdegree > indegree; belonging to SEM: 7, SUB: 3), while in four cases *O. vulgaris* brain areas have the same number of incoming and outgoing.

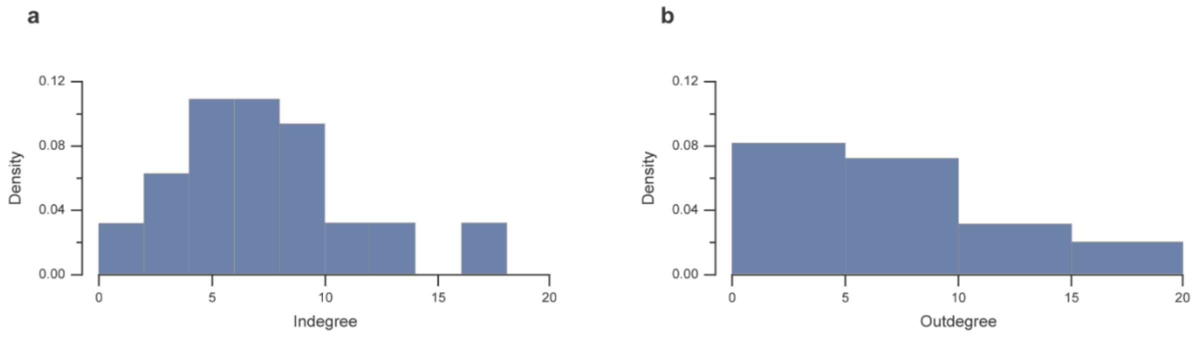

**Supplementary Figure 3**. Distribution of degrees. Probability density of the in-degree (a) and out-degree (b) in the octopus brain network.

### Supplementary Table 5 – Participation coefficients

| gamma 0.6 | | | | | | |
|---|---|---|---|---|---|---|
| **Mass** | **Lobe** | | wm – indeg | wm- outdeg | **P-in** | **P-out** |
| **OL** | *Optic lobe* | **OL** | 0.68 | 1.82 | 0.47 | 0.36 |
| | *Peduncle lobe* | **PEDU** | -0.41 | 1.55 | 0 | 0 |
| | *Olfact lobe* | **OLFL** | -0.77 | 0.23 | 0 | 0 |
| | *Optic gland* | **Optic** | -1.13 | -1.61 | 0 | 0 |
| **SEM** | *Subradular ganglia* | **sRAD** | -1.82 | -1.24 | 0 | 0 |
| | *Inferior buccal ganglia* | **IBL** | -1.82 | -1.85 | 0 | 0 |
| | *Superior buccal lobe* | **SBL** | 0.33 | 0.89 | 0.22 | 0 |
| | *Posterior buccal lobes* | **PBL** | 0.69 | -0.02 | 0 | 0 |
| | *Lateral inferior frontal lobes* | **LatIFL** | 1.41 | 0.59 | 0.17 | 0.40 |
| | *Median inferior frontal lobe* | **mIFL** | -0.02 | -1.24 | 0 | 0.44 |
| | *Subfrontal lobes* | **SUBFL** | -0.38 | -0.32 | 0 | 0 |
| | *Lateral superior frontal lobes* | **SLFL** | 0.69 | -0.32 | 0.32 | 0.28 |
| | *Median superior frontal lobes* | **mSFL** | 0.33 | -0.32 | 0.35 | 0.28 |
| | *Vertical lobes* | **VL** | 0.33 | -0.02 | 0 | 0 |
| | *Subverical lobes* | **SVL** | 1.41 | 0.89 | 0.48 | 0.46 |
| | *Anterior basal lobe* | **ABL** | 0.32 | 0.76 | 0.42 | 0.18 |
| | *Precommissural lobe* | **OC** | -0.77 | -0.03 | 0.50 | 0.38 |
| | *Dorsal basal lobe* | **DBAL** | -0.41 | 0.23 | 0.28 | 0.22 |
| | *Subpedunculate lobe* | **sPED** | -0.04 | -0.56 | 0 | 0.32 |
| | *Median basal lobes* | **mBAL** | 2.49 | 1.55 | 0.36 | 0.32 |
| | *Interbasal lobes* | **IntBAL** | -0.41 | -0.56 | 0.47 | 0 |
| | *Lateral basal lobes* | **LatBAL** | -0.04 | 0.76 | 0.24 | 0 |
| **PER** | *Magnocell lobe* | **MAGNO** | -0.04 | -0.29 | 0.48 | 0.49 |
| **SUB** | *Prebrachial lobes* | **pBRL** | 0.33 | 1.50 | 0.35 | 0.49 |
| | *Postbrachial lobes* | **PBRL** | 0.33 | 1.50 | 0.35 | 0.49 |
| | *Anterior pedal lobes* | **ACL** | -0.41 | -0.82 | 0.47 | 0.48 |
| | *Anterior chromatophore lobes* | **APL** | -1.13 | -1.09 | 0 | 0 |
| | *Lateral pedal lobes* | **LatPL** | 1.77 | -0.29 | 0.26 | 0 |
| | *Posterior pedal lobe* | **PPL** | 1.04 | -0.82 | 0.38 | 0.38 |
| | *Palliovisceral lobe* | **PVL** | -1.10 | 0.59 | 0.50 | 0.49 |
| | *Posterior chromatophore lobes* | **PCL** | -0.77 | -0.82 | 0 | 0 |
| | *Vasomotor lobes* | **VAL** | -0.74 | -0.63 | 0.32 | 0 |

| gamma 1 | | | | | | |
|---|---|---|---|---|---|---|
| **Mass** | **Lobe** | | wm - indeg | wm- outdeg | **P-in** | **P-out** |
| **OL** | *Optic lobe* | **OL** | 0 | 0.5 | 0.71 | 0.64 |
| | *Peduncle lobe* | **PEDU** | 0 | 0.5 | 0.32 | 0.49 |
| | *Olfact lobe* | **OLFL** | 0 | 0.5 | 0 | 0.41 |
| | *Optic gland* | **Optic** | -1.58 | -2 | 0 | 0 |
| **SEM** | *Subradular ganglia* | **sRAD** | -1.59 | -0.87 | 0 | 0 |
| | *Inferior buccal ganglia* | **IBL** | -1.59 | -1.73 | 0 | 0 |
| | *Superior buccal lobe* | **SBL** | 0.53 | 1.30 | 0.53 | 0.35 |
| | *Posterior buccal lobes* | **PBL** | 1.06 | 0 | 0.38 | 0.44 |
| | *Lateral inferior frontal lobes* | **LatIFL** | 0.45 | -1.79 | 0.58 | 0.69 |
| | *Median inferior frontal lobe* | **mIFL** | 0 | -1.30 | 0.44 | 0.67 |
| | *Subfrontal lobes* | **SUBFL** | 0 | 0.43 | 0.32 | 0 |
| | *Lateral superior frontal lobes* | **SLFL** | 0.45 | 0.45 | 0.62 | 0.28 |
| | *Median superior frontal lobes* | **mSFL** | 0.45 | 0.45 | 0.59 | 0.28 |
| | *Vertical lobes* | **VL** | -1.79 | 0.45 | 0.49 | 0.28 |
| | *Subverical lobes* | **SVL** | 0.45 | 0.45 | 0.74 | 0.72 |
| | *Anterior basal lobe* | **ABL** | 0.04 | 0.91 | 0.62 | 0.46 |
| | *Precommissural lobe* | **OC** | -0.80 | 0.47 | 0.72 | 0.41 |
| | *Dorsal basal lobe* | **DBAL** | 0 | 0.5 | 0.5 | 0.53 |
| | *Subpedunculate lobe* | **sPED** | 1.58 | 0 | 0.28 | 0.32 |
| | *Median basal lobes* | **mBAL** | 1.71 | 1.78 | 0.63 | 0.58 |
| | *Interbasal lobes* | **IntBAL** | -0.80 | -0.40 | 0.66 | 0 |
| | *Lateral basal lobes* | **LatBAL** | -0.80 | 1.34 | 0.61 | 0.20 |
| **PER** | *Magnocell lobe* | **MAGNO** | -0.80 | -0.40 | 0.72 | 0.67 |
| **SUB** | *Prebrachial lobes* | **pBRL** | 1.06 | 0.87 | 0.49 | 0.65 |
| | *Postbrachial lobes* | **PBRL** | 1.06 | 0.87 | 0.49 | 0.65 |
| | *Anterior pedal lobes* | **ACL** | 0.04 | -0.83 | 0.53 | 0.48 |
| | *Anterior chromatophore lobes* | **APL** | -0.80 | -1.26 | 0 | 0 |
| | *Lateral pedal lobes* | **LatPL** | 1.71 | 0.04 | 0.47 | 0 |
| | *Posterior pedal lobe* | **PPL** | 0.88 | -0.83 | 0.57 | 0.38 |
| | *Palliovisceral lobe* | **PVL** | -0.53 | 0.43 | 0.5 | 0.64 |
| | *Posterior chromatophore lobes* | **PCL** | -0.38 | -0.83 | 0 | 0 |
| | *Vasomotor lobes* | **VAL** | 0 | 0 | 0.32 | 0 |

| gamma 1.4 | | | | | | |
|---|---|---|---|---|---|---|
| **Mass** | **Lobe** | | wm - indeg | wm- outdeg | **P-in** | **P-out** |
| **OL** | *Optic lobe* | **OL** | 0 | 0.5 | 0.77 | 0.73 |
| | *Peduncle lobe* | **PEDU** | 0 | 0.5 | 0.32 | 0.64 |
| | *Olfact lobe* | **OLFL** | 0 | 0.5 | 0 | 0.45 |
| | *Optic gland* | **Optic** | -1.58 | -2 | 0 | 0 |
| **SEM** | *Subradular ganglia* | **sRAD** | -1.19 | -0.10 | 0 | 0 |
| | *Inferior buccal ganglia* | **IBL** | -1.19 | -1.26 | 0 | 0 |
| | *Superior buccal lobe* | **SBL** | 0.85 | 1.64 | 0.72 | 0.59 |
| | *Posterior buccal lobes* | **PBL** | 0.85 | -0.10 | 0.66 | 0.67 |
| | *Lateral inferior frontal lobes* | **LatIFL** | 0.45 | -1.79 | 0.68 | 0.76 |
| | *Median inferior frontal lobe* | **mIFL** | -0.17 | -0.68 | 0.67 | 0.67 |
| | *Subfrontal lobes* | **SUBFL** | 0.85 | 0.48 | 0.56 | 0.48 |
| | *Lateral superior frontal lobes* | **SLFL** | 0.45 | 0.45 | 0.62 | 0.28 |
| | *Median superior frontal lobes* | **mSFL** | 0.45 | 0.45 | 0.59 | 0.28 |
| | *Vertical lobes* | **VL** | -1.79 | 0.45 | 0.49 | 0.28 |
| | *Subverical lobes* | **SVL** | 0.45 | 0.45 | 0.78 | 0.74 |
| | *Anterior basal lobe* | **ABL** | -0.42 | 1.43 | 0.70 | 0.46 |
| | *Precommissural lobe* | **OC** | -1.01 | 0.80 | 0.72 | 0.41 |
| | *Dorsal basal lobe* | **DBAL** | 0 | 0.5 | 0.50 | 0.56 |
| | *Subpedunculate lobe* | **sPED** | 1.58 | 0 | 0.28 | 0.32 |
| | *Median basal lobes* | **mBAL** | 1.34 | 0.80 | 0.71 | 0.74 |
| | *Interbasal lobes* | **IntBAL** | -1.01 | -0.45 | 0.66 | 0 |
| | *Lateral basal lobes* | **LatBAL** | -1.15 | -0.58 | 0.69 | 0.62 |
| **PER** | *Magnocell lobe* | **MAGNO** | -1.79 | -1.79 | 0.76 | 0.74 |
| **SUB** | *Prebrachial lobes* | **pBRL** | 0.45 | 0.45 | 0.67 | 0.73 |
| | *Postbrachial lobes* | **PBRL** | 0.45 | 0.45 | 0.67 | 0.73 |
| | *Anterior pedal lobes* | **ACL** | -0.42 | -1.07 | 0.66 | 0.48 |
| | *Anterior chromatophore lobes* | **APL** | 0.58 | -0.58 | 0 | 0 |
| | *Lateral pedal lobes* | **LatPL** | 1.34 | -0.45 | 0.63 | 0.32 |
| | *Posterior pedal lobe* | **PPL** | 0.17 | -1.07 | 0.68 | 0.38 |
| | *Palliovisceral lobe* | **PVL** | 0.45 | 0.45 | 0.44 | 0.77 |
| | *Posterior chromatophore lobes* | **PCL** | 0.58 | 1.15 | 0.38 | 0 |
| | *Vasomotor lobes* | **VAL** | 0.45 | 0.45 | 0.32 | 0 |

### Supplementary Figure 4 - Community Structure

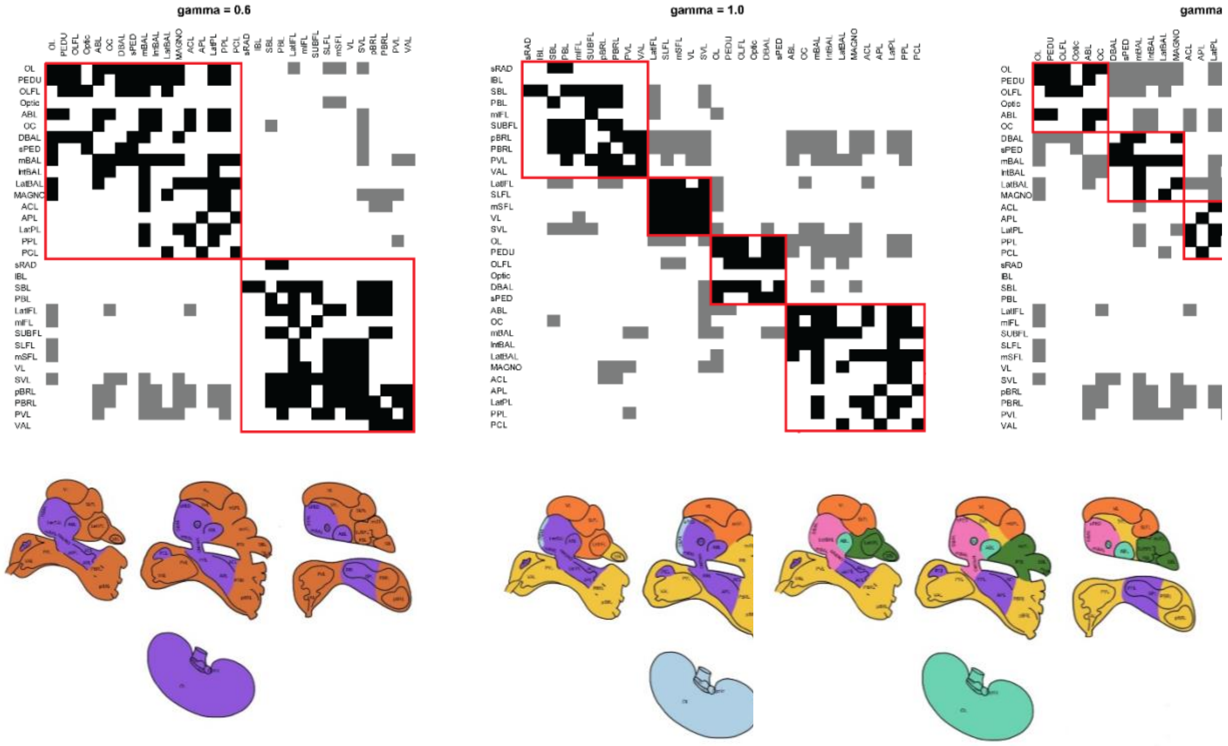

**Supplementary Figure 4** – see legend

**Supplementary Figure 4** (see previous page). Community structure detected in the brain network of *O. vulgaris*. The adjacency matrix is reordered by community assignment (top row); the flatmap projections on sagittal and parasagittal sections of octopus brain and its parts depicting the community assignments (below). See also Figure 3, main text.

With ***gamma = 0.6*** (left panel) two communities were generated (red squares). The first community comprises: optic lobe, peduncle, olfactory, anterior basal, precommissural, dorsal basal, subpedunculate, median basal, interbasal, lateral basal, magnocellular, anterior chromatophore, anterior pedal, lateral pedal, posterior pedal, posterior chromatophore lobes. The second community comprises: subradular, superior buccal, posterior buccal, lateral inferior frontal, median inferior frontal, subfrontal, lateral superior frontal, median superior frontal, vertical, subvertical, pre-brachial, post-brachial, palliovisceral, vasomotor lobes.

Mid Panel: ***γ=1.0*** - four communities are identified (red squares). The first community is represented by: subradular, superior buccal, posterior buccal, median inferior frontal, subfrontal, pre-brachial, post-brachial, palliovisceral, vasomotor lobes. The second by: lateral inferior frontal, lateral superior frontal, median superior frontal, vertical and subvertical lobes. The third including: optic lobe, peduncle, olfactory, dorsal basal and subpedunculate lobes. The fourth one: anterior basal, precommissural, median basal, interbasal, lateral basal, magnocellular, anterior chromatophore, anterior pedal, lateral pedal, posterior pedal, posterior chromatophore lobes.

With ***γ=1.4*** (right panel) six communities were detected (red squares). The first community is represented by: optic lobe, peduncle, olfactory, anterior basal and precommissural lobes. The second by: dorsal basal, subpedunculate, median basal, interbasal, lateral basal and magnocellular lobes. The third including: anterior chromatophore, anterior pedal, lateral pedal, posterior pedal and posterior chromatophore lobes. The fourth community: subradular, superior buccal, posterior buccal, lateral inferior frontal, median inferior frontal and subfrontal lobes. The fifth consisting in: lateral superior frontal, median superior frontal and vertical lobes. The sixth and last one by: subvertical, pre-brachial, post-brachial, palliovisceral and vasomotor lobes.
